# Limiting etioplast gene-expression induces apical hook twisting during skoto-morphogenesis of *Arabidopsis* seedlings

**DOI:** 10.1101/2022.11.02.514823

**Authors:** Salek Ahmed Sajib, Björn Grübler, Cylia Oukacine, Etienne Delannoy, Florence Courtois, Caroline Mauve, Claire Lurin, Bertrand Gakière, Thomas Pfannschmidt, Livia Merendino

## Abstract

When covered by a layer of soil, seedling development follows a dark-specific program (skoto-morphogenesis) consisting of small, non-green cotyledons, a long hypocotyl and an apical hook to protect meristematic cells. We recently highlighted the role played by mitochondria in the high energy-consuming reprogramming of *Arabidopsis* skoto-morphogenesis. Here, the role played by plastids, another energy supplying organelle, in skoto-morphogenesis is investigated. This study was conducted in dark conditions to exclude light signals so as to better focus on those produced by plastids. It was found that limitation of plastid gene-expression (PGE) induced an exaggerated apical hook bending. Inhibition of PGE was obtained at the level of transcription and translation using the antibiotics rifampicin and spectinomycin, respectively, as well as plastid RPOTP RNA polymerase mutants. Rifampicin-treated seedlings also showed expression induction of marker nuclear genes for mitochondrial stress, perturbation of the mitochondrial metabolism, increase of ROS levels and an augmented capacity of oxygen consumption by mitochondrial alternative oxidases (AOX). AOX enzymes act to prevent over-reduction of the mitochondrial electron transport chain. Previously, we reported that AOX1A, the main AOX isoform, was a key component in the developmental response to mitochondrial respiration deficiency. In this work, we suggest the involvement of AOX1A in the response to PGE dysfunction and propose the importance of signalling between plastids and mitochondria. Finally, it was found that seedling architecture reprogramming in response to rifampicin was independent of canonical organelle retrograde pathways and the ethylene signaling pathway.

**Significance statement:** In underground germination conditions, seedling development follows a dark-specific program (skoto-morphogenesis) consisting of small and non-green cotyledons, a long hypocotyl and an apical hook to protect meristematic cells. We show that skoto-morphogenesis is reprogrammed when plastid gene expression is perturbed leading to an exaggeration of apical hook bending. We propose the involvement of the cooperation between plastids and mitochondria, the energy-supplying organelles of the cell.

## Introduction

Plastids are cell organelles in plants that according to the tissue, display large morphological and functional variations^1^. Most of these plastid types can interconvert upon environmental- and/or development-induced changes in plant tissues. For example, in the presence of light, undifferentiated (proplastids) or dark-differentiated (etioplasts) plastids become chloroplasts that are characterized by a complex internal structure of membranes, the thylakoids that house chlorophyll-containing proteins and photosynthetic complexes. The photosynthetic process produces ATP, reducing power and sugars/carbon-skeletons that are supplied to the cell. Etioplasts contain prolamellar bodies and complexes containing protochlorophyllide (Pchlide), NADPH and light-dependent NADPH:Pchlide oxidoreductase. In general, the morphological and functional conversions among plastid types are only possible by changes in plastid proteome composition. The plastid proteome is encoded by both plastid and nuclear genomes. This is due to the fact that during their evolution from symbiotic photosynthetic bacteria to cellular organelles, plastids lost most of the original genetic information that was transferred to the nuclear genome. Through a pathway that is referred to as anterograde, the nucleus codes for plastid proteins and controls plastid physiology. However, plastids contain their own genome and because of their prokaryotic origin, the machinery dedicated to the expression of the plastid genome shares many similarities with the bacterial counterpart and so it is sensitive to antibiotics. In *Arabidopsis thaliana* plants, the plastid genome is transcribed by two different types of RNA polymerases, PEP (Plastid-Encoded RNA Polymerase) and NEP (Nucleus-Encoded RNA Polymerase) in combination with different types of factors^2^. PEP is of the eubacterial-type and it can be blocked by antibiotics such as rifampicin. PEP being the only eubacterial RNA polymerase in the cell, this antibiotic is a highly specific inhibitor of PEP-dependent plastidial transcription. PEP transcriptional activity and specificity are regulated by six nucleus-encoded sigma factors and require the presence of nucleus-encoded PEP Associated Proteins (PAPs). In addition, two NEP RNA polymerases, RPOTmp and RPOTp, participate to plastid genome transcription. RPOTp is exclusively localized in plastids whereas RPOTmp is targeted to both plastids and mitochondria. These phage-type enzymes are insensitive to antibiotics. For the plastid translational machinery, the majority of the components are plastid-encoded and eubacterial-like (70S type-ribosomes) but plastid-specific nucleus-encoded factors are also involved^3^. Plastid translational processes can be blocked by several antibiotics, such as spectinomycin and lincomycin, that specifically affect the 30S and 50S subunits of plastid ribosomes, respectively^4^.

Mitochondria are the site of cellular respiration. Like plastids, they are also semi-autonomous organelles of prokaryotic origin, containing their own genome but depending on the nucleus for the expression of their proteins. Concerning the mitochondrial gene-expression machinery, two NEP RNA polymerases are responsible for transcription of the mitochondrial genome; RPOTm, which is exclusively found in mitochondria, and RPOTmp which is also targeted to plastids. They are phage type enzymes, like RPOTp, and therefore insensitive to antibiotic treatments. On the other hand, mitochondrial ribosomes are of the prokaryotic type.

If the nucleus controls the physiology of both plastids and mitochondria by encoding organelle proteins through anterograde signaling pathways, organelles inform the nucleus about their own state through retrograde signalling pathways. Retrograde signals can be classified as either biogenic or operational^5,6^ according to whether they are generated during organelle biogenesis or from mature organelles, respectively, in response to developmental signals and/or environmental cues. Biogenic signals can be triggered by organelle gene expression. A major effect of plastid retrograde signals is to reduce the expression of PHotosynthesis-Associated Nuclear Genes, PHANGs, including *LHCB1.3* and *RBCS,* whenever chloroplast activity is impaired^7^. On the other hand, mitochondrial retrograde signals induce the expression of Mitochondrial Dysfunction Stimulon (MDS) genes, including *AOX1A, AT12CYS, NDB4* and *UPOX,* when mitochondria become dysfunctional^8^. Key signalling factors responsible for organelle communication with the nucleus belong to the Genome UNcoupled (GUN) class for plastids and to WRKY and NAC families for mitochondria. To date, six GUN factors have been characterized (Susek et al., 1993; Woodson et al., 2011). Among them, only in the case of GUN1, the retrograde signal has been shown to be mediated by plastid gene expression (PGE), whereas GUN2-6 are all connected to the tetrapyrrole biosynthesis pathway. Concerning the mitochondrial retrograde pathways, ANAC017 has been shown to control the transcription of a large set of nuclear MDS genes in response to mitochondrial stress and is therefore considered as a master regulator of this specific signaling pathway^9,10^. However, it is worth mentioning that a wide range of stresses like UV, salt, heat, can induce these genes via other transcriptional networks that do not involve ANAC017/ *mitochondrial* retrograde response (*MRR*) pathways^11^.

During the last decades, many studies have been performed to investigate cross-talk pathways between the nucleus and bioenergetic organelles in light-grown plants. However, in light conditions it is very difficult to separate the effect of plastid signals from the influence of the light as both target the same genes and act on the same promoter sequences. Only limited studies have been performed with dark-grown seedlings but this would allow a better focus on organelle retrograde pathway by excluding “contaminating” light signals.

When seeds are covered by soil, seedlings develop in the dark and they follow a specific developmental program called skoto-morphogenesis^12,13^; exhibiting small, non-green cotyledons, a long hypocotyl and an apical hook that protects meristematic cells during soil emergence. At this early stage of development, cell energy demand is high. Recently, the role played by mitochondria during seedling establishment in the dark was investigated using the *rpoTmp* mutant affected in organelle genome transcription together with other independent mutants defective in cytochrome c oxidase (COX)-dependent respiration^14,15^. This showed that defects in mitochondrial activity led to a drastic re-programming of *Arabidopsis* seedling architecture^16^, characterized by an exaggerated hook curvature and a shortening of the hypocotyl. In addition, *rpoTmp* seedlings activated an alternative oxidase (AOX)-dependent respiratory pathway. AOX, an ubiquinol oxidase, has been shown previously to prevent over-reduction of the respiratory electron transport chain and the formation of harmful reactive oxygen species (ROS)^17^. Genetic impairment of the main AOX isoform (AOX1A) led to perturbations in the redox state of NAD(P)/H pools and ROS contents as well as in the ATP/ADP ratio, thus highlighting the impact of AOX function on cellular energy budget^18,19^. Double mutants affected in RPOTMP and AOX1A enzymes showed that reprogramming of skoto-morphogenesis, and in particular the exaggeration of the apical hook, in response to a respiratory stress was dependent on AOX1A^16^. This study strongly suggested that AOX1A was a key component in the developmental response to mitochondrial dysfunction and that the ANAC017-dependent mitochondrial retrograde pathway was at least partially required for the reprogramming of skoto-morphogenesis.

Even if the role of plastids on photo-morphogenesis has been clearly demonstrated^20^, the impact of plastid function on skoto-morphogenesis has never been investigated. Although non-photosynthetic, etioplasts can still supply energy to the cell through a process called etio-respiration that allows ATP synthesis *via* electron flow from NAD(P)H coming from the oxidative pentose phosphate pathway to oxygen^21^. In addition to energy related processes, etioplasts are also responsible for the biosynthesis of hormones that control development. Dysfunctional etioplast gene expression is therefore expected to impact seedling development even in the dark.

The main objective of this current work was to study the impact of defects in specific steps of PGE, transcription and translation, on nuclear gene-expression and seedling skoto-morphogenesis and to determine the potential role played by organelle retrograde and ethylene signaling pathways in the developmental response.

## Results

### Limitation of plastid transcription and translation alters both skoto- and photo-morphogenesis

To better understand the impact of PGE limitation on dark-grown seedling morphogenesis, PEP-mediated-transcription and translation steps were blocked in plastids by treatment with specific antibiotics, rifampicin and spectinomycin, respectively. A very strong alteration of skoto-morphogenesis, especially at the level of hook bending (increase of the apical hook angle over 180°, identified as a twisting phenotype), was induced by both antibiotics (Fig. 1A). In addition, a similar modification was detected in *rpoTp* seedlings, deficient in the plastid specific and nucleus-encoded RNA polymerase RPOTp^22,23^, suggesting that exaggeration of the hook curvature was not due to pleiotropic effects of the drugs but it was a specific developmental response to plastid gene-expression dysfunction. The twisting phenotype was observed when seeds were submitted to rifampicin treatment either at the beginning of the stratification period, or prior to the exposure time to light required to induce germination or at the beginning of the dark growth phase, thus restricting the effect of the plastid blockage to skoto-morphogenesis *in sensu stricto* (Supplementary Fig. 1). The PGE limitation-induced twisting phenotype was comparable to that brought about by a treatment with ACC, the direct precursor of ethylene (Fig. 1A). Remarkably, impairment of translation by spectinomycin or treatment of seedlings with ACC also strongly reduced the hypocotyl length, conversely to the neutral effect of rifampicin treatment or the lack of RPOTp in the mutant line.

**Fig. 1.**
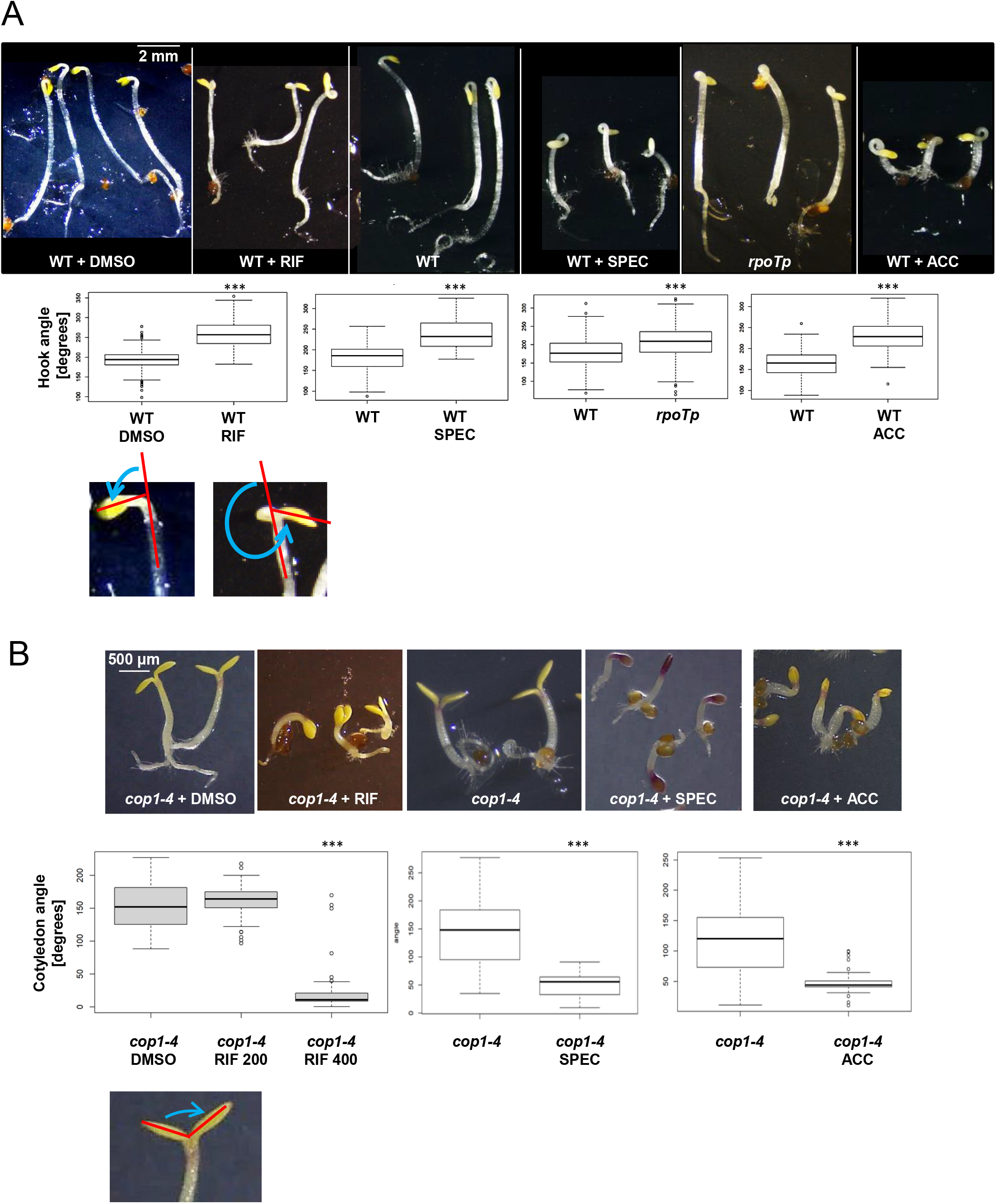
Limitation of PGE interferes with skoto-morphogenesis (A) and photo-morphogenesis (B). **A) Top panel**: Dissection microscope images of etiolated WT seedlings grown in the presence of DMSO (as a mock control for rifampicin), rifampicin (RIF) 200 μg/ml, spectinomycin (SPEC) 250 μg/ml or 1-aminocyclopropane-carboxylic acid (ACC) 20 μM and mutant *rpoTp* seedlings. Scale bar corresponds to 2 mm. **Middle panel**: Box-plots of median values of apical hook angle measurements. The number of pooled individuals (N, measured in 4 independent analyses) corresponds to 137 seedlings for WT + DMSO and 156 for WT + RIF; N (measured in 4 independent analyses) corresponds to 122 seedlings for WT and 135 for WT + SPEC; N (measured in 7 independent analyses) corresponds to 252 seedlings for WT and 246 for *rpoTp*; N (measured in 3 independent analyses) corresponds to 82 seedlings for WT and 115 for WT + ACC. The differences between treated and untreated seedlings or between *rpoTp* seedlings and WT that are significant by statistical tests are indicated by asterisks (***P<0.001). **Bottom panel**: The apical hook curvature was measured as the angle (blue line) that is formed by the two straight lines (red) passing through the hypocotyl and the cotyledon axes. Any hook angle larger than 180° was considered as twisted. **B) Top panel**: Dissection microscope images of etiolated *cop1-4* seedlings grown in the presence of DMSO (as a mock control for rifampicin), rifampicin (400 μg/ml), spectinomycin (500 μg/ml) and ACC (20 μM). Scale bar corresponds to 0.5 mm. **Middle panel**: Box plots of median values of cotyledon separation angle measurements. The number of pooled individuals (N) that were measured in 2 independent cotyledon angle analyses corresponds to 62 seedlings for *cop1-4*+DMSO, 44 for *cop1-4*+RIF 200 and 67 for *cop1-4*+RIF 400; to 74 for both *cop1-4* and *cop1-4*+ACC; the number of pooled individuals (N) that were measured in 3 independent hook angle analyses corresponds to 61 for *cop1-4* and 51 for *cop1-4+SPEC* (500 μg/ml) seedlings. The differences between treated and untreated seedlings that are significant by statistical tests are indicated by asterisks (***P<.001). **Bottom panel**: The cotyledon separation angle was measured as the angle (blue line) that is formed by two straight lines (red) passing through the cotyledon axes.

We also tested the effect of PGE limitation on photo-morphogenesis. Being rifampicin light-sensitive, the impact of PGE inhibition was tested in the dark on the development of constitutive photo-morphogenic mutant *cop1-4* seedlings (short hypocotyl, absence of hook, and cotyledon opening even in the dark (Fig. 1B, see *cop1-4*+DMSO, *cop1-4)).* Measurements of cotyledon opening angles indicated that PGE limitation by rifampicin or spectinomycin as well as ACC treatment impaired cotyledon opening thus suppressing the *cop1* mutant seedling phenotype. A similar interference with photo-morphogenesis has been described already for lincomycin- (another antibiotic targeting plastid translation) and ACC-treated light grown seedlings^24^.

### Limitation of plastid transcription impacts on the expression of nuclear genes linked with mitochondrial biology and on mitochondrial metabolism

To determine the reprogramming extent of gene expression brought about by the RIF-induced limitation of plastid transcription, we performed an Affymetrix microarray analysis to obtain a whole transcriptomic profiling of etiolated WT seedlings grown in the presence of rifampicin *versus* control conditions. Only 51 genes (41 nuclear and 10 mitochondrial) genes showed a log2-fold change (fc) >1.5 with a p-value lower than 0.05; 43 nuclear genes showed a log2-fc <-1.5 with a p-value lower than 0.05 (Table S1). These data indicated a restricted impact of the RIF-induced limitation of plastid transcription on nuclear gene-expression. Functional grouping of the most up- and down-deregulated genes according to the GO term revealed a strong link with mitochondrial biology (organization, protein import, respiration chain) and to stress/hormone response (ethylene, acid abscisic, Supplementary Fig. 2A and B). The ten most up-regulated genes can be clustered in distinct classes: one group containing markers of mitochondrial stress/targets of the mitochondrial retrograde pathway (*AT12CYS-2* and its co-expression partner *NDB4* ^25^, three *DUF295 Organellar B* genes^26^ and *SPL2* phospatase^27^) and a gene involved in general mitochondrial functions, *TOM7-2* (coding for a member of the family of TOM7 translocases of the outer mitochondrial membrane); a gene coding for a chloroplast-localized protein with an oxido-reduction function; a group included genes encoding two proline- and glycine-rich proteins, often involved in stress response (Fig. 2A). Among these ten genes, seven encoded mitochondrial-localized proteins and eight were among the ten most up-regulated genes in the mitochondrial respiration mutant *rpoTmp*^16^. In addition, mitochondrial genes were globally over-expressed, probably as a compensation effect for a mitochondrial stress (Fig. 2B, p-value=2.2e-16). On the other hand, as expected for a plastid specific inhibitor of transcription, expression of plastid genes was globally down-regulated (Fig. 2C, p-value=0.000188). The impact of the rifampicin treatment was comparable on the expression of the different cellular genomes as indicated by the similar range of the log2FC (Fig. 2B-D).

**Fig. 2.**
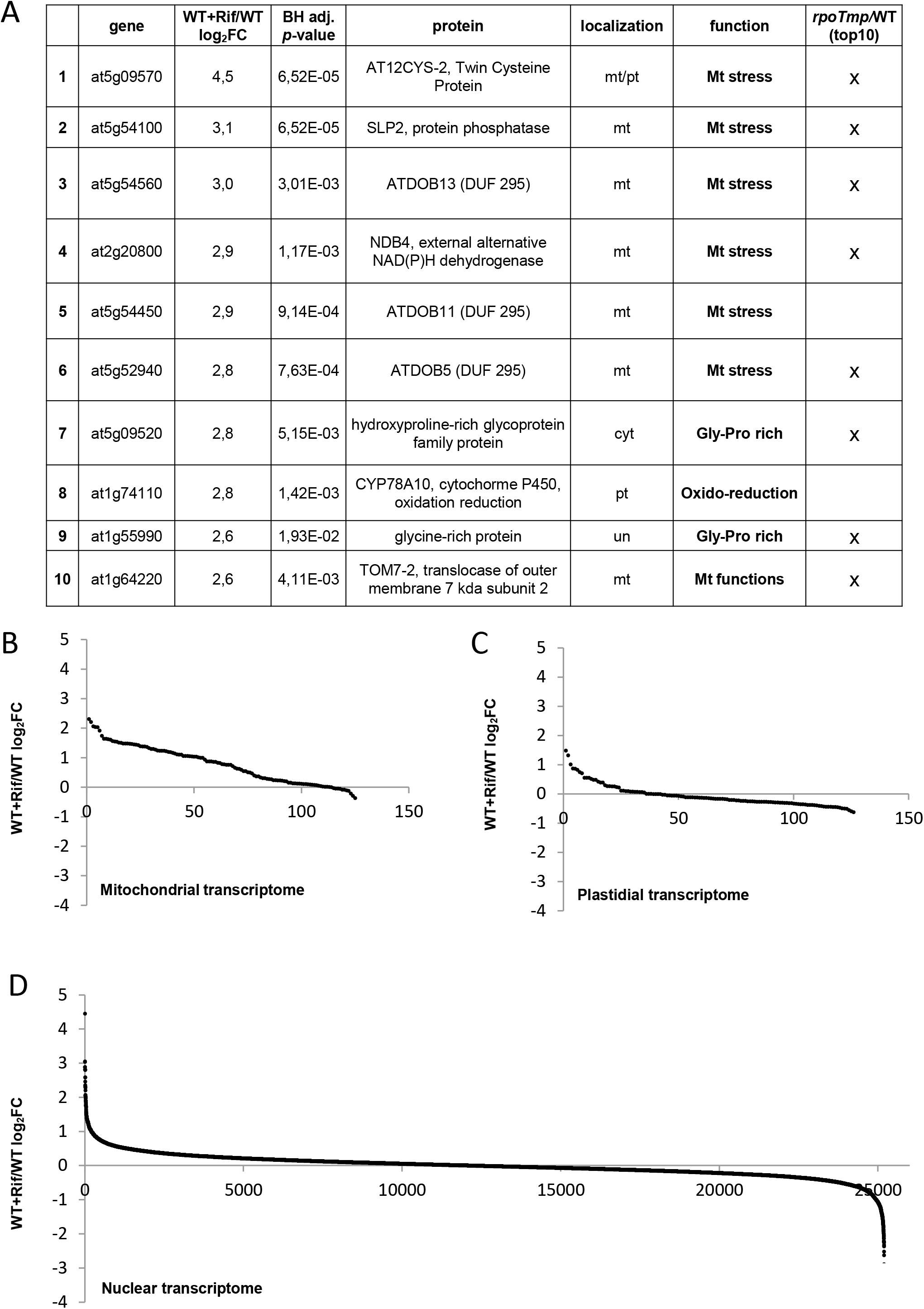
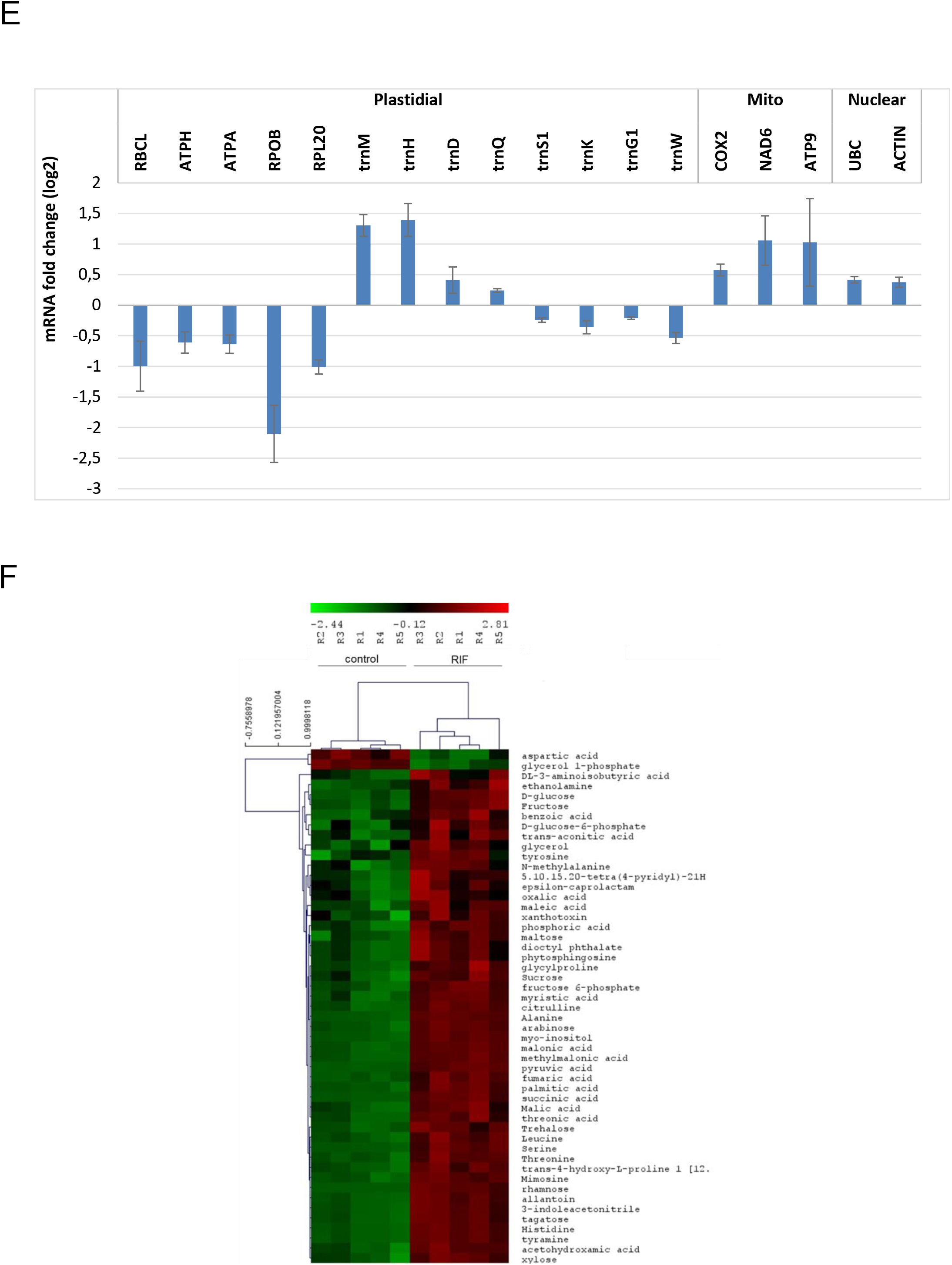
Relative gene expression and metabolic profiles of rifampicin-treated *versus* control WT etiolated seedlings. **A)** Relative expression values for the 10 most up-regulated genes in rifampicin-treated etiolated WT seedlings are presented as log2-fold changes (FC) together with the corresponding p-values, which are derived from a t-test adjusted for false discovery rate (FDR) after the Benjamini-Hochberg (BH) procedure (BH adj. *p*-value). Gene and protein identities are indicated together with the cellular localization (mitochondrial (mt), plastidial (pt), cytoplasmic (cyt), uncharacterized (un)) and function (“Mt stress” stands for “response to mitochondrial stress”). On the right side, it is indicated when the genes are also found among the most ten up-regulated genes in etiolated mutant *rpoTmp* seedlings^16^. **B)** Relative expression values for mitochondrial, **C)** plastidial and **D)** nuclear genes in rifampicin-treated etiolated WT seedlings. Given values represent log2-fold changes (FC). **E**) Log2 fold change (FC) of expression levels of plastidial, mitochondrial and nuclear genes in RIF-treated WT seedlings *versus* untreated seedlings determined by RT-qPCR. The expression levels were normalized to the mean of *PP2A* (nuclear) expression, used as a reference gene. The mean values of two biological replicates are plotted. Error bars correspond to standard errors. **F**) Metabolic Heat Map of control (DMSO) and rifampicin-treated seedlings (RIF). In total, the levels of 51 metabolites were significantly affected by RIF treatment (T-test, p-value 0.05); the amounts of 49 metabolites were increased while two metabolites decreased. Metabolomic profiling of five repetitions (R1-5) was performed using Gas Chromatography-Mass Spectrometry (GC-MS).

RT-qPCR analyses were also performed with independent biological replicates and we could confirm the decrease of some of the plastid mRNAs analyzed in rifampicin-treated seedlings, as already observed in the microarray analysis (Fig. 2E). The levels of transcripts that are generally approved as PEP-dependent as *atpH, atpA* and *rbcL,* and NEP-dependent as *rpoB* were all impacted by the rifampicin treatment that is PEP-specific. In addition, as already indicated by the microarray data, some of the plastid tRNAs that are generally agreed as PEP-dependent were over-accumulated upon rifampicin treatment, either because over-expressed by the NEP machinery, as compensatory mechanism, or for increased stability. The classification of PEP and NEP-dependent genes based on studies performed in light-grown plants is likely not valid in our dark conditions. Finally, RT-qPCR data indicate that the levels of both mitochondrial and nuclear transcripts are either increased or unchanged by the rifampicin treatment supporting that this antibiotic limits specifically plastid transcription.

We also tested the specificity of spectinomycin and found that the levels of the mitochondrial-encoded protein NAD9 were only slightly decreased by the treatment with this antibiotic, while the signals corresponding to the plastid-encoded S7 protein are almost undetectable (Suppl Fig. 3). These data indicate that spectinomycin blocks specifically plastid translation when supplied to dark grown seedlings.

In addition to the induction of nuclear genes related to mitochondrial biology, RIF treatment is also responsible for metabolic perturbation and in particular for upregulation of intermediates of the mitochondrial TCA process. Metabolic profiling by GC-MS revealed 51 significantly affected metabolites in RIF-treated seedling *versus* control (T-test, p-value 0.05), 49 metabolites increased in abundance and two metabolites decreased (Fig. 2F, Suppl Fig. 4, Table S3). All glycolytic-related carbohydrates (e.g., sucrose, glucose, fructose, rhamnose, xylose, and maltose) as well as TCA cycle intermediates (including citrate, malate, fumarate, and succinate) were significantly more abundant in the PGE-limited seedlings. This could sign a respiratory defect in relation to an over-reduction of the pyridine nucleotide pool. Supporting this, we found an accumulation in lactate that is as well a signature of mitochondrial respiratory deficiency and a shift to fermentation, as already observed in *rpoTmp* and *atphb3* mutants^10^. As mitochondrial perturbation can lead to less ATP synthesis, especially in non-photosynthetic tissues, altering amino-acids levels, we quantified these nitrogen metabolites using OPA-HPLC. Even though the total amino acid content was unchanged, we observed a strong shift in amino-acid metabolism, with decreased amounts of Asn, Lys, Gln that might be a consequence of lack of energy (ATP) required for their synthesis. On the other hand, the pyruvate-derived branched chain amino acids Ala, Leu, Val, Ser and Trp were accumulated, likely as a consequence of the accumulation of the glycolytic intermediates. And the Gly levels were reduced, likely because of a defect in the mitochondrial process producing Gly from Ser. Furthermore, a substantial increase in fatty acids (palmitic acid, stearic acid, and malonic acid) was observed in the RIF-treated seedlings. These data point to mitochondrial perturbation in the PGE-limited seedlings.

### The organellar retrograde pathways that are dependent on either ANAC017 or GUN1 are not involved in the developmental response to PGE dysfunction

In order to determine if the ANAC017-dependent mitochondrial retrograde signalling pathway was involved in the observed developmental response to antibiotic-mediated PGE limitation, *anac017* mutant seedlings were dark-grown on rifampicin. RT-qPCR analyses showed that genetic impairment of ANAC017-dependent retrograde signalling pathways did not interfere with the nuclear gene expression response (induction of mitochondrial stress marker genes, Fig. 3A). In addition, *anac017* seedlings still showed the developmental response (twisting phenotype) to rifampicin-induced plastid transcriptional limitation (Fig. 3B). To analyze the involvement of the plastid retrograde signalling pathway, *gun1* seedlings were dark-grown on rifampicin and seedling morphogenesis was analyzed (Fig. 3C). Like *anac017* plants, *gun1* seedlings responded at the developmental level (hook twisting) to the rifampicin treatment. We then determined the impact of the rifampicin treatment on the expression of the Photosynthetic Associated Nuclear Genes (PhANGs) that are responsive to the plastid retrograde signaling pathway in light-grown plants. To achieve this, we used the transcriptomic profiling data of etiolated WT seedlings grown in the presence/absence of rifampicin and found that none of the selected genes was differentially expressed in a statistically significant manner (Supplementary Fig. 5A, BH adj P-value > 0.05).

**Fig. 3.**
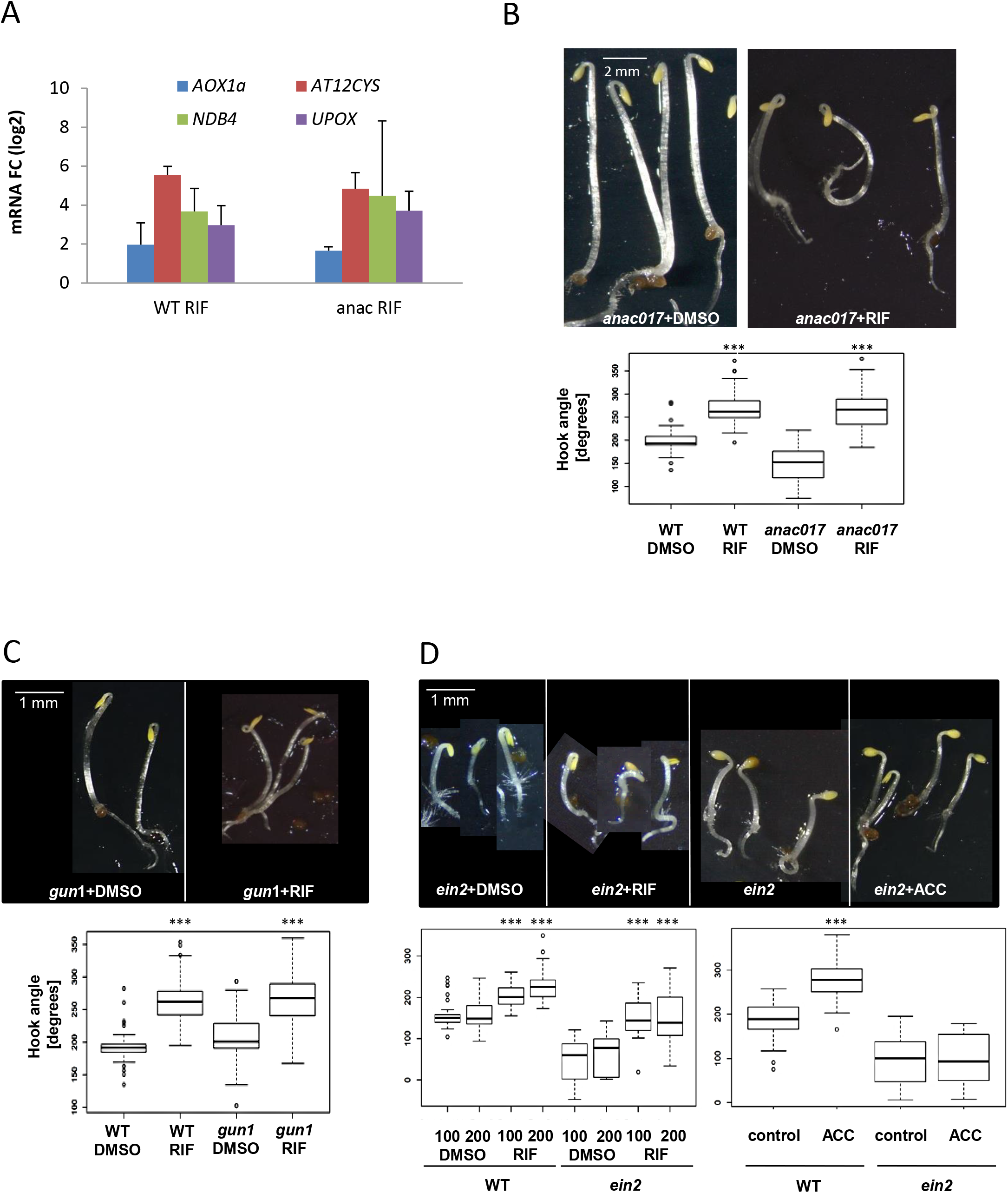
Control of hook bending under PGE limitation is independent of the ANAC017- and GUN1-dependent organelle retrograde pathways and of the EIN2-dependent signalling pathway. **A)** Log2 fold change (FC) induction of expression levels of mitochondrial stress marker genes in RIF-treated WT and mutant *anac017* seedlings *versus* corresponding untreated seedlings determined by RT-qPCR. The expression levels were normalized to the mean of *PP2A* expression, used as a reference gene. The mean values of two biological replicates are plotted. Error bars correspond to standard errors. **B) Top panel:** Dissection microscope images of etiolated *anac017* seedlings grown in the presence of DMSO as a mock control for rifampicin and rifampicin (RIF) 200 μg/ml. **Bottom panel:** Box plots of median values of apical hook angle measurements. The number of pooled individuals (N) that were measured in 2 independent hook angle analyses corresponds to 40 seedlings for WT+DMSO and 44 for WT+RIF; to 46 for *anac017*+DMSO and 54 for *anac017*+RIF. The differences between treated and untreated seedlings that are significant by statistical tests are indicated by asterisks (***P<.001) **C) Top panel**: Dissection microscope images of etiolated *gun1-201* seedlings grown in the presence of DMSO or rifampicin (RIF) 200 μg/ml. **Bottom panel**: Box plots of median values of apical hook angle measurements. The number of pooled individuals (N) measured in 2 independent hook angle analyses corresponds to 40 seedlings for WT+DMSO and 41 for WT+RIF; to 54 for *gun1-201+DMSO* and 46 for *gun1-201+RIF.* The differences between treated and untreated seedlings that are significant by statistical tests are indicated by asterisks (***P<.001) **D) Top panel**: Dissection microscope images of etiolated *ein2-1* seedlings grown in the presence of DMSO, rifampicin (RIF) 200 μg/ml or ACC 20 μM. **Bottom panel**: Box plots of median values of apical hook angle measurements. The number of pooled individuals (N) that were measured in 2 independent hook angle analyses corresponds to 36 seedlings for WT + DMSO and 37 for WT + RIF (100 μg/ml); to 43 for *ein2-1* + DMSO and 29 for *ein2-1* + RIF (100 μg/ml); N (measured in 2 independent analyses) corresponds to 30 seedlings for WT + DMSO and 42 for WT + RIF (200 μg/ml); to 50 for *ein2-1* + DMSO and 62 for *ein2-1* + RIF (200 μg/ml). N (measured in 2 independent analyses) corresponds to 73 seedlings for WT and 79 for WT + ACC; to 33 for *ein2-1* and 36 for *ein2-1* + ACC. The differences between treated and untreated seedlings that are significant by statistical tests are indicated by asterisks (***P<.001).

### The EIN2-dependent ethylene signalling pathway is not involved in the developmental response to PGE dysfunction

The twisting phenotype of seedlings induced by treatment with rifampicin or ACC was comparable, suggesting an eventual role of the ethylene-dependent signaling pathway in the response to the PGE limitation. To test this hypothesis, we observed the skoto-morphogenic profile of ethylene-insensitive mutant *ein2-1* seedlings in the presence of rifampicin. It was found that they still exhibited a twisting phenotype when dark-grown on rifampicin containing medium, whereas the exaggeration of hook bending was no longer detected in the presence of ACC (Fig. 3D). In addition, using the transcriptomic data, none of the selected genes involved in ethylene biosynthesis and signaling were induced in a statistically significant manner (Supplementary Fig. 5B, BH adj p-value> 0.05) except for ACS4 that showed a log2 fold change of 2 and the BH adj p-value inferior to 0.05. However, no differences were found in ethylene emission levels when rifampicin-treated seedlings were compared to mock (DMSO) seedlings (Supplementary Fig. 5C). In conclusion, such results exclude the involvement of the ANAC017, GUN1 and EIN2-dependent pathways in the observed developmental response to PGE dysfunction.

### Genetic disruption of AOX1A impairs the developmental response to PGE limitation

AOX1A has been shown to control skoto-morphogenesis in response to mitochondrial respiration deficiency^16^. Here we observed that limitation of PGE by rifampicin induced an increase of AOX gene expression not only at the transcript level (Fig. 3A) but also especially at the protein level (Fig. 4A). High levels of AOX proteins also corresponded to an increase in the AOX capacity (KCN-insensitive respiration, Fig. 4B) but not to the total respiratory O_2_ consumption rate in rifampicin-treated seedlings (Supplementary Fig. 6). In order to investigate if AOX played a role in the developmental response to PGE limitation, two independent allelic mutants in the *AOX1A* gene, coding for the major AOX isoform in dark-grown seedlings^16^, *aox1a-1* and *aox1a-2,* were treated with rifampicin. 40-50% of the mutant seedling population was characterized by the absence of twisted hooks and presented either hook or “comma” phenotype (Fig 4C and Supplementary Fig. 7A). The “comma” phenotype is arbitrary defined by the absence of sharp hook bending. Importantly, a much higher percentage of seedlings with hook and comma phenotypes, 70%, was detected when the rifampicin concentration was increased to the double, RIF 400μg/ml. When RIF-treated *aox1a* seedlings presenting twisted, hook or comma phenotypes were exposed to light, they greened (Supplementary Fig. 7B). Finally, we performed NBT staining to detect the accumulation of ROS and found that the RIF-induced superoxide levels were highly induced by genetic abolishment of AOX1A (Fig. 4D). These data highlight a functional link between PGE limitation, the presence of AOXa1, ROS accumulation and the twisting response.

**Fig.4.**
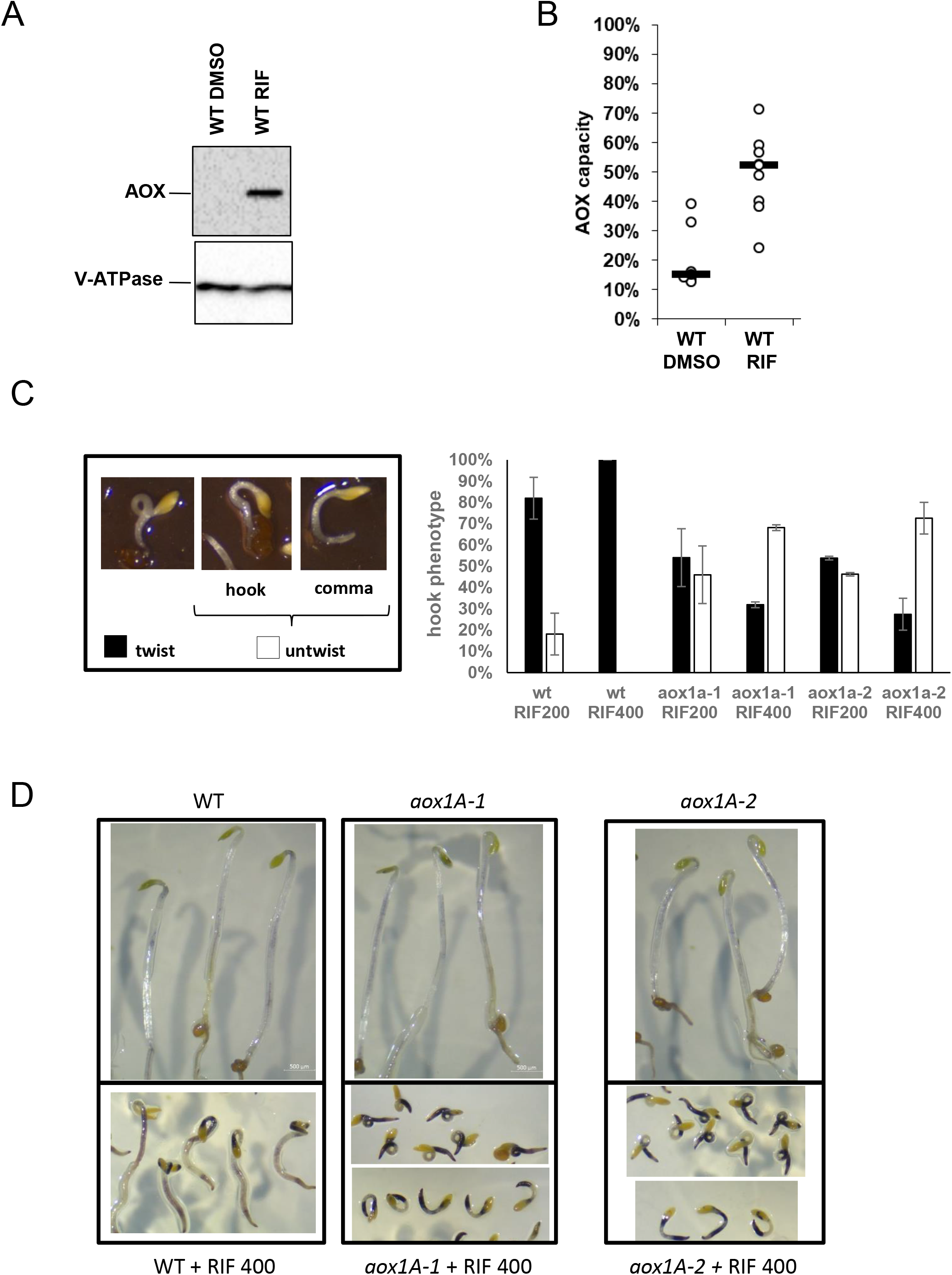
Impact of AOX1A genetic abolishment on the apical hook bending under plastid gene-expression inhibitory conditions. **A)** Analysis of AOX1A protein content used as a marker for mitochondrial stress. Protein extracts (20μg) from etiolated WT seedlings grown in the presence of DMSO or rifampicin (RIF) 200 μg/ml were separated by SDS-PAGE and immunoblotted with specific antisera against mitochondrial AOX protein (nuclear-encoded) and the vacuolar protein ATPase (nuclear-encoded) as a loading control. **B)** Capacity of the AOX-dependent pathway in etiolated WT seedlings grown in the absence or presence of rifampicin (RIF) 200 μg/ml. AOX capacity was calculated upon addition of 1 mM (final concentration) KCN to the measurement cell and shown as the % ratio of KCN-insensitive O_2_ consumption rate to total O_2_ consumption rate. Median values (thick horizontal lines) of N independent measurements (full circles) were scatter-plotted for WT (N=7) and WT+RIF (N=9). **C)** Phenotypic analysis of RIF-treated *aox1a* mutant seedlings. **Left panel**: Zoomed dissection microscope images of RIF (200 μg/ml)-treated *aox1a-1* seedlings showing twist and untwist phenotypes. The untwisted class comprises seedlings with hooks that are either sharp (hooked) or round (comma). **Right panel:** Percentage of the WT, *aox1a-1* and *aox1a-2* seedlings grown on rifampicin (200 and 400 μg/ml) and showing either twist or untwisted phenotype. The number of pooled individuals (N) measured in 2 independent hook angle analyses corresponds to 83 for WT+RIF200, 132 for WT+RIF400, 69 for *aox1a-1*+RIF200, 160 for *aox1a-1*+RIF400, 120 for *aox1a-2*+RIF200 and 164 for *aox1a-2*+RIF400 seedlings. **D)** Microscope images of NBT-stained WT, *aox1a*-1 and *aox1a-2* seedlings, that were grown in control conditions or in presence of RIF 400.

## Discussion

Here it is shown that the early developmental program of dark-grown Arabidopsis seedlings is altered when PGE is limited in the presence of either rifampicin or spectinomycin (Figure 1A). Rifampicin is highly specific to the plastid transcription machinery since it targets PEP, the only prokaryotic RNA polymerase in a plant cell^28^. Microarray and RT-qPCR analyses were performed in this work showing the decrease of many plastid transcripts analyzed in rifampicin-treated seedlings (Figs 2C and E, Table S1). We have also observed that the levels of mitochondrial transcripts are increased and of nuclear transcripts unchanged by the rifampicin treatment and we conclude that rifampicin is specific to plastid transcription (Figs 2B, D and E, Table S1). Spectinomycin is an antibiotic that targets the plastid ribosomal machinery. However, since the mitochondrial ribosomal machinery is also of the prokaryotic type, it might also be inhibited by this drug. However, the levels of the mitochondrial-encoded protein NAD9 were only slightly decreased by the treatment with spectinomycin (Suppl Fig. 3), while the ones of the plastid-encoded S7 protein are almost undetectable. Our data indicate that spectinomycin blocks specifically plastid translation when supplied to dark grown seedlings. Treatment of seedlings with either of the two antibiotics tested induced the exaggeration of the apical hook angle (Figure 1A). However, shortening of the hypocotyl was observed only in the case of spectinomycin. Considering that spectinomycin targets translation of all plastid transcripts while rifampicin inhibits transcription of only the PEP polymerase-dependent plastid genes, the degree of impact on skoto-morphogenesis (only hook over-bending or an additional impact on hypocotyl length) might be related to the degree of plastid gene-expression limitation. The conclusion that the twisting phenotype was specifically induced by blockage of PGE was strongly supported by the observation that apical hooks were over-bended in seedlings mutated in the plastid nucleus-encoded RNA polymerase RPOTp (Figure 1A). Also, in *rpoTp* mutant seedlings transcription of only a subset of plastid genes, in this specific case driven by RPOTp-dependent promoters, was inhibited and the impact was detected exclusively at the hook bending level and not at the hypocotyl length.

The remark that the plastid dysfunction-promoted twisting phenotype was comparable to the ACC-induced apical hook over-bending suggested the involvement of ethylene. However, treatment with rifampicin still affected the architecture of ethylene insensitive *ein-2* mutant seedlings, while in this mutant no effect was detected upon ACC treatment (Figure 3D). In addition, no increase in the accumulation of ethylene (Supplementary Figure 5C) and transcripts of ethylene responsive genes (apart from ACS4, Supplementary Figure 5B) could be detected in 3-day-old etiolated WT seedlings treated with rifampicin. Finally, the other traits of the ethylene-induced triple response observed in WT seedlings treated with ACC such as hypocotyl thickening and root shortening were not observed in rifampicin-treated seedlings (Figure 1A). All of these data indicate that the ethylene-dependent signaling pathway is not involved in the developmental reprogramming of skoto-morphogenesis observed in response to rifampicin-induced PGE dysfunction in dark-growth conditions.

It was also found that limitation of PGE by rifampicin and spectinomycin treatments impaired cotyledon separation in the constitutive photo-morphogenic mutant *cop1* in dark-growth conditions (Figure 1B). Recently, it has been shown that lincomycin, another antibiotic targeting plastid translation, impaired cotyledon separation during early development of light-grown WT seedlings^20^. Taken together, these data highlight the impact of limiting both plastid transcription and translation on seedling development even when considering different light regime growth conditions.

To better understand the molecular signalling mechanisms underlying the developmental response to PGE limitation in the dark, the transcriptomic profile of rifampicin-treated seedlings was analyzed using microarrays (Table S1) and it was found that six of the top ten most up-regulated transcripts were markers for mitochondrial stress (Figure 2A). In particular, *AT12CYS-2* (At5g09570) was found to be induced mostly in mitochondrial stress condition^25,29^. In addition, RT-qPCR and western analyses showed that the expression of AOX1A, another marker gene for mitochondrial stress, was clearly induced at both the transcript and protein levels (Fig. 3A and 4A). The stimulation of mitochondrial stress markers was surprising for an antibiotic treatment targeting PGE (Fig. 2C, plastidial transcriptome was generally down-regulated, and Fig. 2E) and not mitochondrial gene expression (Fig. 2B, mitochondrial transcriptome was overall induced, and Fig. 2E). However, recent reports have indicated that the expression of marker genes for mitochondrial stress such as *AOX1A* was also induced in response to plastidial stress, as in the case of plants that are treated with methyl viologen, a molecule generating ROS in chloroplasts ^30^ and etiolated *immutans* seedlings lacking plastid terminal oxidase PTOX, an enzyme involved in etio-respiration that is responsible for ATP synthesis in dark-grown seedlings^21^. These data suggest the existence of cross-talk signaling pathways between organelles and the nucleus. However, the rifampicin-driven twisting phenotype was observed even when ANAC017-dependent mitochondrial retrograde pathways were genetically impaired (Figure 3B). Furthermore, the induction of mitochondrial stress marker genes *AOX1A*, *At12CYS-2* and *NDB4* by rifampicin treatment was not altered by the absence of ANAC017 (Fig. 3A). One interpretation of the data is that an alternative mitochondrial retrograde pathway, independent of canonical factors, is being used when seedlings are subjected to mitochondrial stress in response to plastid dysfunction in the dark. Alternatively, we cannot exclude that induction of stress marker gene expression and of the developmental response does not go *via* a mitochondrial route.

Metabolomics profiling was performed in rifampicin-treated WT Arabidopsis seedlings to determine which metabolites are associated with the PGE limitation and seedling growth (Fig. 2F and Supplementary Fig. 4). GC-MS analysis revealed an increase in a number of TCA and glycolytic intermediates in rifampicin-treated seedlings, supporting the idea that rifampicin treatment impacts on mitochondrial metabolism. Previous studies have demonstrated that in mitochondrial mutants free amino acids accumulate due to a lack of cellular energy^10^. Here, we observed a strong shift in amino-acids metabolism, probably due to a lack of ATP production not compensated by photosynthesis, impairing Lys, Asn and strongly Gln synthesis. These data support further that RIF interferes with energy production. However, global amino-acid pool remained unchanged. Among shikimate-derived amino-acids, only the anthranilate-branch of the pathway seems to be upregulated, as shown by an increase of Trp levels, and could impact auxin synthesis involved in apical hook formation. Earlier report stated that in the case of mitochondrial dysfunction mutants (*atphb3, rpoTmp),* several other metabolites such as fatty acids and lactic acid are altered^10^. Here, we also noticed the accumulation of various fatty acids in the rifampicin-treated seedlings, such as palmitic acid and stearic acid. Specially, lactic acid seems to accumulate, which agrees to the previous report of slowing of aerobic respiration metabolism and a shift to fermentation, which is a classical sign of limited mitochondrial function. Finally, the decrease in Gly can be interpreted as a defect in the mitochondrial process producing Gly from Ser.

We have also investigated whether the plastid retrograde pathway was involved in the signalling behind the expression and developmental response to PGE limitation. The rifampicin-induced twisting phenotype was observed even when the GUN1 retrograde pathway was genetically impaired (Fig. 3C). Furthermore, the expression levels of canonical gene targets of the plastid retrograde pathway remained unchanged in WT seedling upon rifampicin treatment (Supplementary Fig. 5A, BH adj p-value>0.05). Therefore, the limitation of plastid transcription by rifampicin does not induce a plastid GUN1-dependent retrograde response in etiolated seedlings. However, the levels of GUN1 protein have not been analyzed in the dark so we cannot certify that GUN1 is accumulated and/or active in darkness. In addition, since the expression levels of PhANG genes were low in the dark^31^, we cannot exclude that an eventual decrease due to the rifampicin treatment might be difficult to detect. As already reported in the case of mitochondrial stress, accumulation of AOX protein was also induced, together with a concomitant increase of its capacity (measured as KCN-insensitive respiration) in response to plastid dysfunction (Figures 4A and B). Importantly, genetic abolishment of the major AOX1A isoform in the dark partially impaired the developmental response to PGE limitation as seen by a sub-set of rifampicin 200-treated *aox1a-1* and *aox1a-2* seedlings, around 40%, being characterized by the absence of a twisted apical hook and presenting either a normal hook (hook) or a round hook (comma, Fig. 4C and Supplementary Fig. 7A). When the concentration of the RIF was increased to 400 μg/ml, the percentage of seedlings that were unable to twist was larger, around 70%. Interestingly, a similar comma phenotype has been observed previously in *aox1a-1/rpoTmp* double mutant seedlings^16^. How AOX1A acts to modify seedling architecture during skoto-morphogenesis is still unclear. AOX is an enzyme involved in the control of 1) ROS and nitric oxide (NO) homeostasis by preventing over-reduction of the mitochondrial electron transport chain^17^ and 2) the redox state of NAD(P)/H pools and ATP/ADP balance, thus influencing cell energy budget^18,19^. We have performed NBT staining to detect ROS accumulation in etiolated seedlings (Fig. 4D). We found that the RIF treatment leads to high ROS levels, especially when AOX1A was genetically abolished. We propose that the ROS produced because of the plastid/mitochondrial stress together with the ones coming from the seeds might inhibit the twisting response. In addition to the involvement in the control of seedling morphogenesis, AOX1A also participates to breaking seed dormancy through the regulation of ROS levels^32^. These concordant results reveal a new role for AOX1A in controlling the early steps of plant development^33^. However, we cannot exclude that the alteration of the mitochondrial status is just a side effect of the rifampicin treatment and that there is not a functional link between mitochondrial perturbation, AOX1A-controlled ROS accumulation and developmental response of the seedlings.

In conclusion, limitation of PGE by rifampicin and spectinomycin leads to skoto-morphogenesis reprogramming especially at the apical hook bending (Fig. 5). PGE limitation is somehow perceived by the nucleus activating the expression of nuclear genes that are considered as markers for mitochondrial stress such as *AOX1A*. This leads to increased mitochondrial AOX protein amounts and AOX-dependent respiration capacity. In RIF-treated *aox1a* mutant seedlings, we observed a strong accumulation of ROS and the incapacity of a sub-population of seedlings to respond with the twisting phenotype. These data support the involvement of mitochondrial AOX1A in controlling seedling architecture through ROS regulation in a situation of plastid/mitochondrial dysfunction. Finally, rifampicin treatment was also shown to modify the accumulation of mitochondrial metabolic intermediates. Communication from plastids to the nucleus could occur directly or *via* the mitochondria using signals passing between the organelles, potentially through physical connections. However, regulation of nuclear expression and the developmental response to PGE limitation are independent of GUN1-dependent plastid and ANAC017-dependent mitochondrial retrograde signalling pathways. Finally, the ethylene signalling pathway was shown not to be involved in the skoto-morphogenic reprogramming of the apical hook upon treatment with plastid specific antibiotics.

**Fig. 5.**
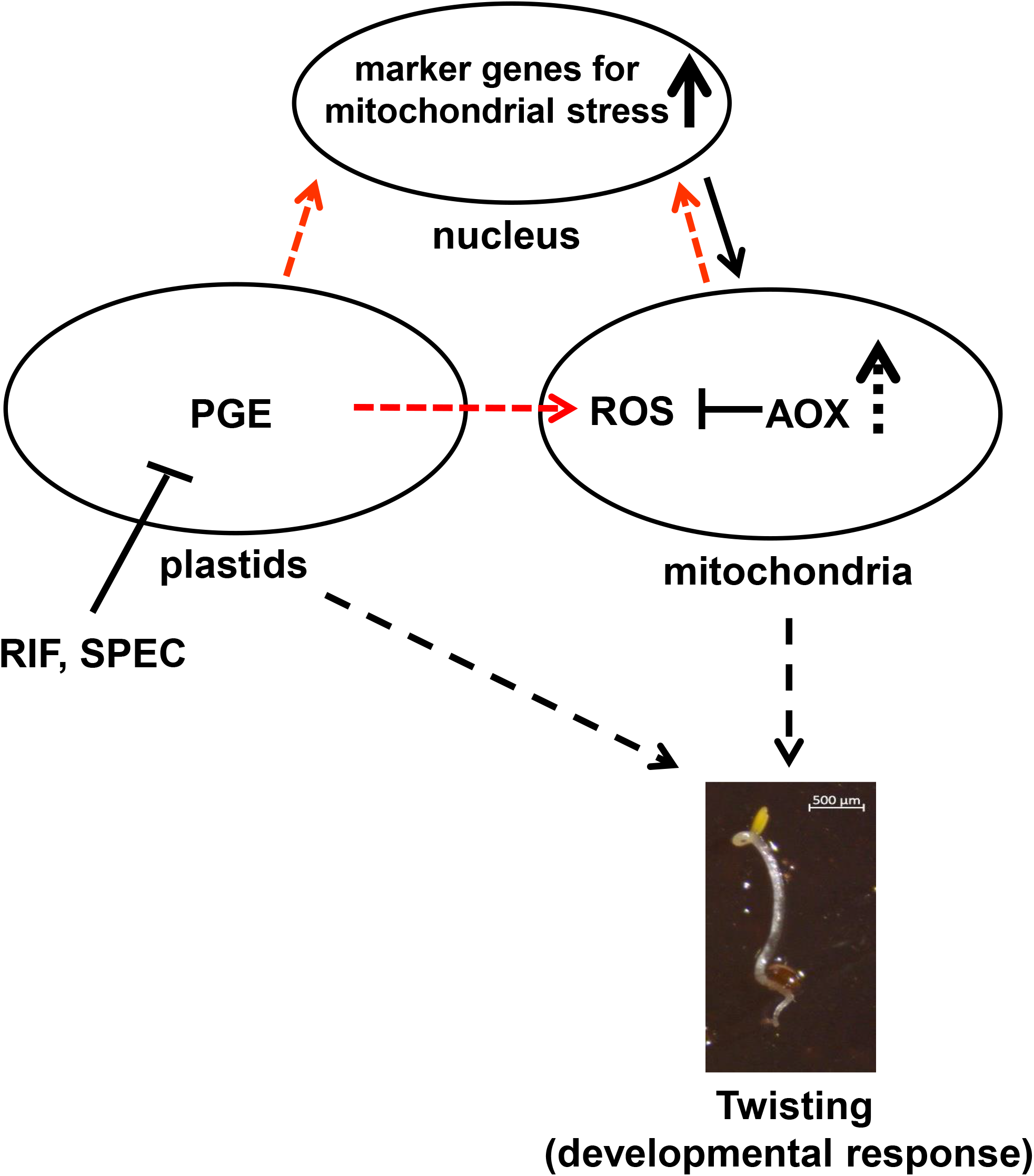
Scheme of the proposed functional link between PGE limitation and skoto-morphogenic reprogramming. In etiolated rifampicin- or spectinomycin-treated WT seedlings, transcription of the PEP-dependent plastid genome or translation of plastid transcripts is limited, respectively. The plastidial stress leads to transcriptional induction of nuclear genes that are considered as markers for mitochondrial stress (thick black arrow, *AOX1A*, *AT12CYS*, *NDB4* and *UPOX*). If nuclear gene-expression is directly regulated by plastid dysfunction or *via* mitochondrial stress is still unknown (red arrows). The activation of nuclear gene expression is independent of both the GUN1-plastidial and the ANAC017-mitochondrial retrograde pathways and might be supported by yet unknown factors. Expression of AOX is increased also at the protein level (thick discontinuous black arrow) and consequently, the capacity of the AOX-dependent chain is upregulated and more electrons can be diverted by this pathway. Role of AOX consists in detoxifying the cell from ROS. As an ultimate effect, a developmental response (twisting phenotype) is induced. If and how the developmental response is triggered directly by plastid signals or *via* mitochondria by AOX-generated signals is still unknown (thin discontinuous black arrows, online version in color).

## Material and Methods

### Plant material

*Arabidopsis thaliana* wildtype (WT) (ecotype Columbia) and mutant lines (*cop1-4*^34^; *rpoTp* (*SALK_067191*^22,23^); *ein2-1* (*CS3071*^35^); *gun1-201* (*SAIL_290_D09*^24^); *anac017* (*SALK_022174, rao2-1*)^6,7^; *aox1A-1* (SALK_084897) and *aox1A-2 (SAIL*_030_D08)^36^) were used in this study. Seeds were surface-sterilized and sown on Murashige and Skoog (MS) agar plates supplemented with 1% sucrose and 0.08% charcoal (as a powder to increase image background of etiolated plants, SIGMA-ALDRICH). For chemical treatments, MS agar plates were supplemented with rifampicin (stock of 150 mg/ml in DMSO; SIGMA-ALDRICH R3501; final solution of 100, 200 and 400 μg/ml, it binds to bacterial-like RNA polymerase); spectinomycin (SIGMA-ALDRICH S4014; final solution of 250 or 500 μg/ml, it binds to the ribosomal 30S subunit); 1-aminocyclopropane-carboxylic acid (ACC, SIGMA-ALDRICH A3903, final solution of 20 μM, it is the direct precursor of ethylene). DMSO-containing MS agar plates were used as a mock-control for the rifampicin treatment.

Seeds were stratified for 3 days (at 4°C in the dark), exposed to light (100 μEm^-2^s^-1^ white light) for 6 h and then grown at 23°C in the dark. Plants were either observed or harvested after 3 days.

In the experiment relative to Supplementary Fig. 1, seeds were sown on nitrocellulose membrane filters (Whatman 10407970, 0.45 μm) either directly on rifampicin-containing MS agar plates at the beginning of stratification or on MS agar plates and then transferred (on the filter) to rifampicin-containing MS agar plates before or after light exposure.

For microarray, RT-qPCR and western-immuno-blot analyses, etiolated plants were harvested in the dark under a green safe-light, immediately frozen in liquid nitrogen, ground in a mortar and subjected to further techniques as described below.

### Measurements of phenotypic parameters

Phenotypic appearance of plantlets was analyzed as previously described^16^. Briefly, digital images of individual plants were taken with a dissection microscope (Olympus SZX12) using the ACT-1C for DXM1200C software. The angles formed in the apical hook between the hypocotyl and cotyledons were subsequently determined using the ImageJ program (Fig. 1A, lower panel). Any hook angle larger than 180° was considered as twisted. For *cop1-4* mutant seedlings, to determine the cotyledon separation, the angle was measured between cotyledons (Fig. 1B, lower panel).

### Statistical analyses of phenotypic parameter measurements

All experimental dataset were tested for the ANOVA assumption of normality of the residuals by both a Normality plot of the residuals and the Shapiro-Wilk test on the ANOVA residuals. Considering that the Shapiro-Wilk test failed, a Wilcoxon test was performed. Statistical analysis was performed using R 4.1.2^37^.

In Fig 1A all comparisons were significantly different (ANOVA p-values<2e-16). In Fig 1B, the cop mutant was insensitive to RIF200 (Wicoxon p-value >0.05) but impacted by RIF400 (ANOVA p-value <1.7e-11), SPEC (ANOVA p-value <2e-16) and ACC (Wilcoxon p-value<2e-16). In Fig 3B for *anac017,* the response to RIF is even higher than in WT (2-way ANOVA p-value = 7.34e-5). In Fig 3C the *gun1* mutant was not significantly different from WT in both DMSO and RIF treatments (Wilcoxon p-values >0.05). In Fig 3D for *ein2,* the response to RIF 100 or 200 is even higher than in WT for RIF 100 and RIF 200 (2-way ANOVA p-value = 0.0003 and 0.006). No response to ACC was observed in *ein2* as opposed to WT (2-way ANOVA p-value =1.29e-11).

### Microarray analysis

Total RNA purification and microarray analysis were performed as indicated in ^16,31^. Total RNA was isolated using RNeasy Plant Mini Kit (Qiagen) with on-column DNase I (Qiagen) treatment. Affymetrix whole transcriptome microarray analysis was performed on three biological replicates by the commercial service Kompetenz-Zentrum für Fluoreszente Bioanalytik (KFB) (Regensburg, Germany). cDNAs were prepared using the Ambion® Whole Transcriptome (WT) Expression Kit and fragmented and labeled using the Affymetrix GeneChip® WT Terminal Labeling Kit. Expression analyses were performed using the “GeneChip® Arabidopsis Gene 1.0 ST Array”. Since the “Ambion® WT Expression Kit” uses a mixture of oligo-dT and random hexamer primers for the generation of the first strand cDNA, the resulting hybridization signals reflect the accumulation of both organellar and nuclear transcripts.

### Statistical analysis of microarray data

Analysis of microarray data was performed as indicated in^16,31^. The cell files containing the scanned chips received from KFB were analyzed with RobiNA^38^ with the background correction method RMA (robust multi-array average expression measure)^39^, p-value correction method Benjamini-Hochberg^40^ and the analysis Strategy Limma for quality control^41^. For the main analysis, RMA normalization with p-value correction using the BH method and a “nestedF” multiple testing strategy was chosen. In the analysis using Excel (MS-Office), data with the description “Multiple Hits” were ignored and only results with one-to-one correspondence to a given gene identity were kept for further analysis (Table S1).

In Fig 2 the distributions of the log2FC of the plastid, mitochondrial and nuclear transcripts were compared using a Wilcoxon test to find global variations between each pool of transcripts. As the transcriptomic normalisation step is applied globally to all genes and that nuclear transcripts are the vast majority of the genes, if the distribution of the organellar transcripts (either plastid or mitochondrial) differs from the distribution of nuclear transcripts then plastid or mitochondrial transcription can be considered to be mis-regulated. In Fig 2B the log2 FC of mitochondrial transcripts in seedlings treated with RIF were globally higher than nuclear transcripts in Fig. 2D (Wilcoxon p-value <2e-16). In Fig 2C the log2 FC of plastidial transcripts in seedlings treated with RIF were globally lower than nuclear transcripts in Fig. 2D (Wilcoxon p-value = 0.000188).

### RT-qPCR analysis of RNA levels

Ground plant material was resuspended in 3 volumes of solution A (10 mM Tris-HCl pH 8.0; 100 mM NaCl; 1 mM EDTA; 1% SDS) and 2 volumes of phenol/chloroform/isoamyl alcohol (25:24:1; v/v/v). After centrifugation, RNAs in the aqueous phase were again extracted twice with phenol-chloroform and finally once with chloroform. After over-night precipitation in 2 M LiCl at 4°C, RNAs were precipitated in ethanol, washed in ethanol 70% and resuspended in water. 500 ng of DNAse (Max Kit, Qiagen) treated-RNAs were retro-transcribed using random hexo-nucleotides or reverse gene-specific primers and the Super-Script II enzyme (Invitrogen) according to the manufacturer’s protocol. The PCR reaction was performed with a Biorad CFX384TM Real Time System PCR machine and forward and reverse gene-specific primers (0.5 μM, Table S2) using the SYBR® Premix Ex Taq™ (Tli RNaseH Plus) from Takara Bio. Data were analyzed using the CFX Manager Software. The expression levels of the transcripts of interest were normalized to the levels of *PP2A* expression, used as a reference gene. The mean values of biological and/or technical replicates were plotted together with the error bars corresponding to standard errors.

### Metabolic analyses

Col-0 Arabidopsis seedlings grown in the presence of 200 μg/ml rifampicin and DMSO control were utilized for metabolic profiling using GC-MS. Ground fresh samples (50 mg FW) were resuspended in 1 mL of frozen (−20°C) water:acetonitrile:isopropanol (2:3:3) containing Ribitol at 4 μg mL^-1^, shaken at 4°C for 10 minutes, centrifuged and 100μl of the SN were dried for 4 hours at 35°C in a Speed-Vac before storing at −80°C. All GC-MS analysis steps were executed as detailed previously (Fiehn, 2006; Fiehn et al., 2008).

For **GC-MS profiling,** the samples were dried again in a Speed-Vac evaporator for 1.5 hours at 35°C before adding 10 μL of 20 mg mL^-1^ methoxyamine in pyridine to the samples and incubating for 90 minutes at 30°C with continuous shaking. Then, 90 μL of N-methyl-N-trimethylsilyl-trifluoroacetamide (MSTFA) (Regis Technologies, USA) were added and incubated for 30 minutes at 37°C. Then, 100μl samples were transferred to an Agilent vial for injection after cooling. 1μl of sample was injected in split-less mode on an Agilent 7890B gas chromatograph linked to an Agilent 5977A mass spectrometer after 4 hours derivatization where the column was an Rxi-5SilMS from Restek (30 m with 10 m Integra-Guard column).

For saturated chemical quantification, an injection in split mode with a ratio of 1:30 was conducted systematically. The oven temperature ramp was 60°C for 1 minute, followed by 10°C min^-1^ to 325°C for 10 minutes. The constant flow of helium was 1.1 mL min^-1^. Temperatures were as follows: Injector temperature was 250°C, transfer line temperature was 290°C, source temperature was 230°C, and quadrupole temperature was 150°C. After a 5.90-minute solvent delay, the quadrupole mass spectrometer was turned on and scanned from 50 to 600 m/z. Absolute retention times were locked to the internal standard d27-myristic acid using the RTL system provided in Agilent’s Masshunter software. Retention time locking decreases the run-to-run variability of retention time. The samples were randomized. In the center of the queue, a mixture of fatty acid methyl esters (C8, C9, C10, C12, C14, C16, C18, C20, C22, C24, C26, C28, C30) was injected for external RI calibration. AMDIS (http://chemdata.nist.gov/mass-spc/amdis/) was used to analyze raw Agilent data files. For metabolite identifications, the Agilent Fiehn GC/MS Metabolomics RTL Library (version June 2008) was used, which is the most comprehensive library of metabolites, comprising GC/MS spectra and retention times for about 700 common metabolites. Peak areas were obtained in splitless and split 30 modes using the Mashunter Quantitative Analysis (Agilent Technologies, CA, USA). Since automatic peak integration can occasionally produce errors, integration was manually validated for each chemical in all analyses. For comparison, the resulting areas were compiled into a single MS Excel file. Ribitol and Dry Weight were used to normalize peak regions. The contents of metabolites were represented in arbitrary units (semi-quantitative determination). All examinations were carried out independently five times. Normalized data (mean-center) were generated into a clustered metabolomic array (heat map) for metabolomics using MeV 4.1 open-source software (Saeed et al., 2003). Metabolites that differed substantially between the two groups were identified using the Student’s T-test and a significance value of P≤ 0.05.

For amino acids quantification using **OPA-HPLC profiling,** the 200 μL of each sample was dried again in a Speed-Vac evaporator for 1.5 hours at 35°C followed by addition of 1.3 ml H_2_O and filtration to inject into the HPLC. The OPA reagent was made 48 h before first use by dissolving OPA at 10mg/mL in 200 μL of methanol and adding 1.8 mL of sodium borate 0.5M (pH 9.5) and 40 μL of 2-mercaptoethanol. The reagent was filtered into an autosampler vial and used for up to 3 days. Precolumn derivatization was performed in the injection loop by automated mixing of 10 μL sample and 10 μL OPA reagent, followed by a delay of 2 min prior to injection. The HPLC (Alliance Waters 2695, Waters Corporation, USA) fitted with a fluo detector (Multi λ Fluorescence Detector 2475) was used. Compounds were measured at λ excitation of 340nm and λ emission of 455 nm. The chromatographic separation was performed with Symmetry C18 column (3.5μm, 4.6*150 mm) by gradient elution at 40 °C using buffer A (20% methanol, 80% sodium acetate, 1% tetrahydrofuran, pH 5.9) and buffer B (80% methanol, 20% sodium acetate, pH 5.9). Buffer flow rate was 0.8 mL/min throughout and total run time per injection was 42 min. The chromatography data were analyzed by Empower software. Peak identity was confirmed by co-elution with authentic standards. Concentration was calculated with calibration curve using peak area of compound of interest.

### Western blot analysis

Total protein extracts were prepared by re-suspending 100 mg of ground plant material in 100 μl of protein lysis buffer (50 mM Tris pH 8.0; 2% SDS; 10 mM EDTA; protease inhibitors (Roche (04693159001) cOmplete™, Mini, EDTA-free Protease Inhibitor Cocktail)) and incubated at room temperature for 30 min. Extracts were cleared by centrifugation for 30 min at 12000 × g at 4°C and dilutions of the supernatant were used to quantify protein amounts with Bradford reagent (SIGMA-ALDRICH B6916). Samples were then denatured for 5 min at 95 °C in 4X reducing electrophoresis sample buffer (200 mM Tris pH 6.8, 5% mercaptoethanol, 4% SDS, 0.2% Bromo Phenol Blue, 20 % glycerol). Comparable amounts of plant protein extracts (100 μg) were separated by SDS-PAGE (12 % acrylamide), electro-blotted onto a nylon membrane that was milk saturated and immuno-decorated with specific polyclonal rabbit antisera against plant mitochondrial alternative oxidase 1 and 2 (AOX1/2; Agrisera, AS04 054), mitochondrial-encoded NAD9^42^, plastidial-encoded S7 (Agrisera, AS15 2877) and vacuole V-ATPase (Agrisera, AS07 213). The blots were developed with Clarity Western ECL substrate (Bio-Rad, Hercules, USA). Images of the blots were obtained using a CCD imager (Chemidoc MP, Bio-Rad) and the Image Lab program (Bio-Rad, Hercules, USA).

### In planta O_2_ consumption measurements

Etiolated plantlets were harvested from Petri dishes and immediately soaked in the dark with 1 mL of oxygenated phosphate buffer (10 mM sodium phosphate buffer pH 7.2; 10 mM KCl; 10 mM glucose) in the measurement cell of a Clark oxygen electrode (Hansatech Instruments). For respiratory electron transfer inhibition experiments, KCN (0.1 M stock, inhibitor of COX-dependent pathway at the level of complex IV) and SHAM (0.1 M stock in DMSO, inhibitor of the AOX-dependent pathway) were added directly into the electrode chamber to a final concentration of 1 mM, once the recorded signal reached a constant value. The capacity (or maximum activity) of the AOX-dependent pathway (corresponding to the SHAM-sensitive/KCN-insensitive pathway) and of the COX-dependent pathway (corresponding to the SHAM-insensitive/KCN-sensitive pathway) was calculated as ratio of O_2_ consumption rate upon addition of either KCN or SHAM and total O_2_ consumption rate (measured in the absence of inhibitors), respectively. The extra mitochondrial O_2_ consumption was evaluated at the end of each measurement after addition of both KCN and SHAM and systematically subtracted from O_2_ consumption rates.

### NBT staining of ROS

Etiolated seedlings were transferred in 12 well plate, vacuum-infiltrated in the dark with 2.5 ml NBT 6 mM staining solution for 5 minutes and incubated for 4h in dark at room temperature with continuous shaking. Digital images of individual plants were taken with a dissection microscope (Olympus SZX12) using the ACT-1C for DXM1200C software.

### Ethylene measurements

For ethylene measurements, etiolated seedlings (fresh weight: 500 mg) were collected and put into 5 ml amber, flat bottom screw neck vials (MACHEREY-NAGEL GmbH & Co. KG, Germany Cat no. 702293), and sealed with an N 9 PP screw cap with a center hole and silicone white/PTFE red septum (MACHEREY-NAGEL GmbH & Co. KG, Germany, Cat no. 702287.1). After 4 h, ethylene accumulation measurements were performed with the ETD-300 ET detector (Sensor Sense B.V., Nijmegen, The Netherlands). The accumulated ethylene was drawn through a valve controller over a period of 7 min and with a constant flow of 3L/h and sent into the laser acoustic spectrometer/detector where ethylene was specifically detected. Levels of ethylene emissions are given in ppbv (parts per billion by volume) as a function of the relative time since the start of the experiment.

## Acknowledgements

This work was supported by the LabEx Saclay Plant Sciences-SPS (ANR-10-LABX-0040-SPS) to IPS2; grants from the Deutsche Forschungsgemeinschaft [PF323-5-2] and the DFG research group FOR 804; the Centre National de la Recherche Scientifique [PEPS] to TP; the French Ministry of Education and the Grenoble Alliance for Integrated Structural Cell Biology (LabEx GRAL, ANR-10-LABX-49-01) to LPCV. We thank Michael Hodges and Emmanuelle Issakidis-Bourguet from IPS2 (Orsay, France) for helpful discussions; Géraldine Bonnard (IBMP, Strasburg, France) for sharing NAD9 antisera. We thank Olivier Van Aken (Lund University, Sweden) for sharing *anac017* seeds, Fredy Barneche (IBENS, France) for *gun1-201* and *cop1-4* seeds and Kristina Kühn (Universität Halle, Germany) for *aox1a* seeds. RNA sample processing and Affymetrix microarray hybridization were carried out at the genomics core facility: Center of Excellence for Fluorescent Bioanalytics (KFB, University of Regensburg, Germany). SAS was supported by fellowship from the Ministère de l’Enseignement supérieur, de la Recherche et de l’Innovation (MESRI) of French Government (Doctoral School of Plant Sciences (SEVE), Université Paris-Saclay) for his Ph.D.

## Author contribution

LM and TP designed research; SAS, BGr, CO, FC, CM and LM performed experiments, ED performed statistical analyses, CL contributed analytical tools. SAS, BGr, FC, CM, BGa, TP and LM analyzed data, LM wrote the manuscript with the help of all co-authors. All authors read and approved the manuscript.

---

**Supplementary Figure 1: The effect of PGE limitation is restricted to skoto-morphogenesis *in sensu stricto.* A)** Etiolated WT seedlings were grown on nitrocellulose membrane filters either on MS agar plates in the absence of rifampicin (A) or on rifampicin-containing MS agar plates since the beginning of stratification (B) or on MS agar plates and then transferred (on the filter) to rifampicin-containing MS agar plates before light exposure (C) or at the beginning of dark-growth (D). The continuous black line indicates the length of the RIF treatment during the growth protocol. **B)** The percentage of seedlings exhibiting hook bending higher than 180° (twist) in a single experiment is reported.

**Supplementary Figure 2: Gene ontology sub-grouping of deregulated genes (DEG) in rifampicin-treated *versus* untreated dark-grown WT seedlings.** Only genes with relative expression values- (log2 fold change) >1.5 **(A)** or <-1.5 **(B)** and p-values<0.05 (Table S1) are given.

**Supplementary Figure 3: Analysis of plastid specificity for the spectinomycin antibiotic.** Analysis of S7 and NAD9 protein contents used as plastidial and mitochondrial markers, respectively. Protein extracts (100μg) from etiolated WT seedlings grown in absence or in presence of spectinomycin (Spec) 250 and 500 μg/ml were separated by SDS-PAGE and immunoblotted with specific antisera against plastidial-encoded S7 and mitochondrial-encoded NAD9 proteins and the vacuolar protein ATPase (nuclear-encoded) as a loading control.

**Supplementary Figure 4: Schematic representation of metabolite ratio by GC-MS and amino acid quantification by OPA-HPLC in untreated and RIF-treated WT seedlings. A)** In case of metabolite ratios, units are arbitrary, while for amino acids levels the units are pmol/mg FW. Statistical significance of differences in metabolite abundance between control and RIF-treated seedlings was verified using the Student’s T-test and a significance value of P≤ 0.05. Asterisks stand for P<0.05(*), P<0.01(**), P<0.001(***). **B)** Total amino acid levels are represented for control and RIF-treated seedlings.

**Supplementary Figure 5: Relative expression values for A) PhANGs and B) genes involved in ethylene biosynthesis and signalling pathways in rifampicin-treated versus untreated etiolated WT seedlings.** Values are given as log2-fold changes (FC) together with the corresponding p-values, which are derived from a t-test adjusted for false discovery rate (FDR) after the Benjamini-Hochberg (BH) procedure (BH adj p-value). Gene and protein identities and function (synthesis or signaling) are also indicated. SAM: S-adenosyl methionine synthase; ACS: ACC synthase; ACO: ACC oxidase. **C)** Ethylene emission levels from 500 mg (fresh weight) of WT etiolated seedlings grown either in the presence of DMSO as a mock control for rifampicin or with rifampicin (RIF) 200 μg/ml. The plot shows the measured ethylene levels in ppbv (parts per billion) during a relative time since the start of the experiment.

**Supplementary Figure 6: Total O_2_ consumption rate in etiolated WT plants grown in the presence or absence of rifampicin.** Median values (thick horizontal lines) of N independent measurements (full circles) were scatter-plotted for mock (N=7) and rifampicin (200 μg/ml)-treated (N=9) WT seedlings.

**Supplementary Figure 7: Dissection microscope images of etiolated *aox1a-1* and *aox1a-2* seedlings. A)** Allelic mutant seedlings, *aox1a-1* and *aox1a-2*, were grown either in the presence of DMSO as a mock control for rifampicin or with rifampicin (RIF) 200 μg/ml. RIF-treated *aox1a* mutant seedlings show twist, hook or comma phenotypes (Fig. 4C). **B)** Dissection microscope images of *aox1a-2* mutant seedlings that were grown on RIF400 for 4 days in the dark or **C)** for 10 days in the dark and then transferred 8 days in the light. Seedlings present comma (1, 2) and twist (3) phenotypes.

**Table S1: Microarray-based relative gene expression profiling (total, plastidial and mitochondrial) of rifampicin-treated *versus* untreated etiolated WT seedlings.** Values are ordered from the highest to the lowest. Microarray-based relative gene expression profiling of etiolated *rpoTmp* versus WT seedlings are also reported for comparison.

**Table S2: Primers used in qPCR expression analyses.**

**Table S3: GC-MS relative metabolomic profiling and amino acid quantification using OPA-HPLC profiling in rifampicin-treated *versus* untreated etiolated WT seedlings.**

