## Supplementary figures and tables for "Limiting etioplast gene-expression induces apical hook twisting during skoto-morphogenesis of *Arabidopsis* seedlings": Supp fig final.pdf

A

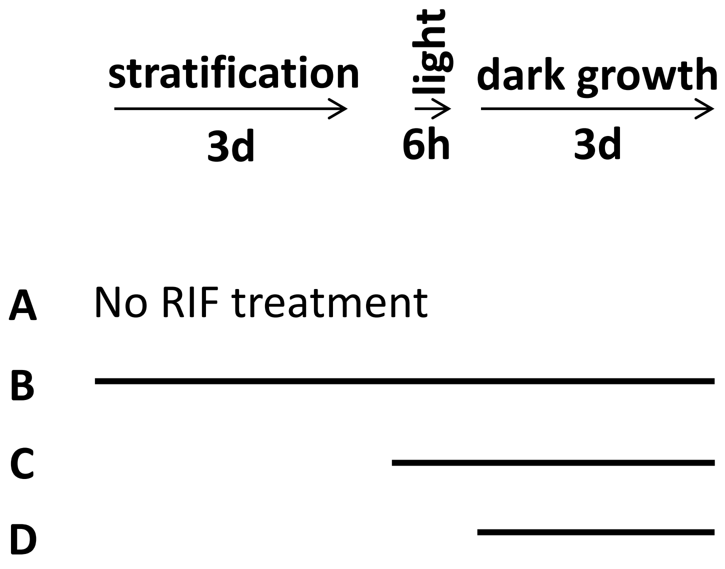


B

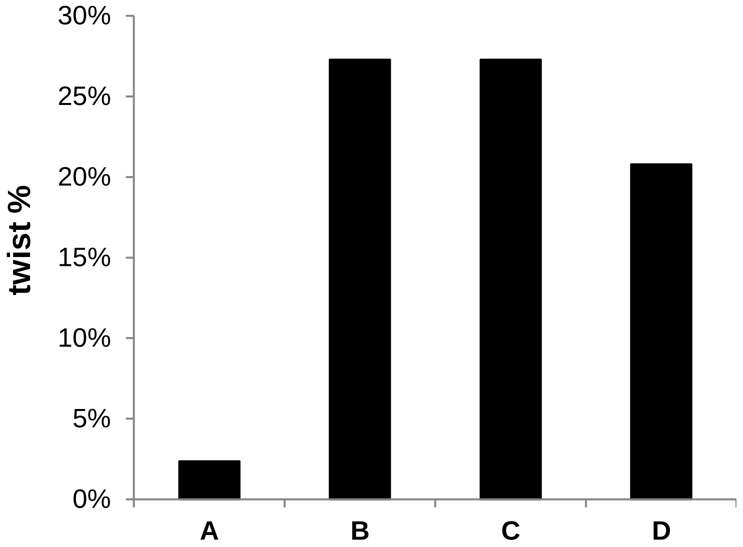


Suppl Fig. 1

A

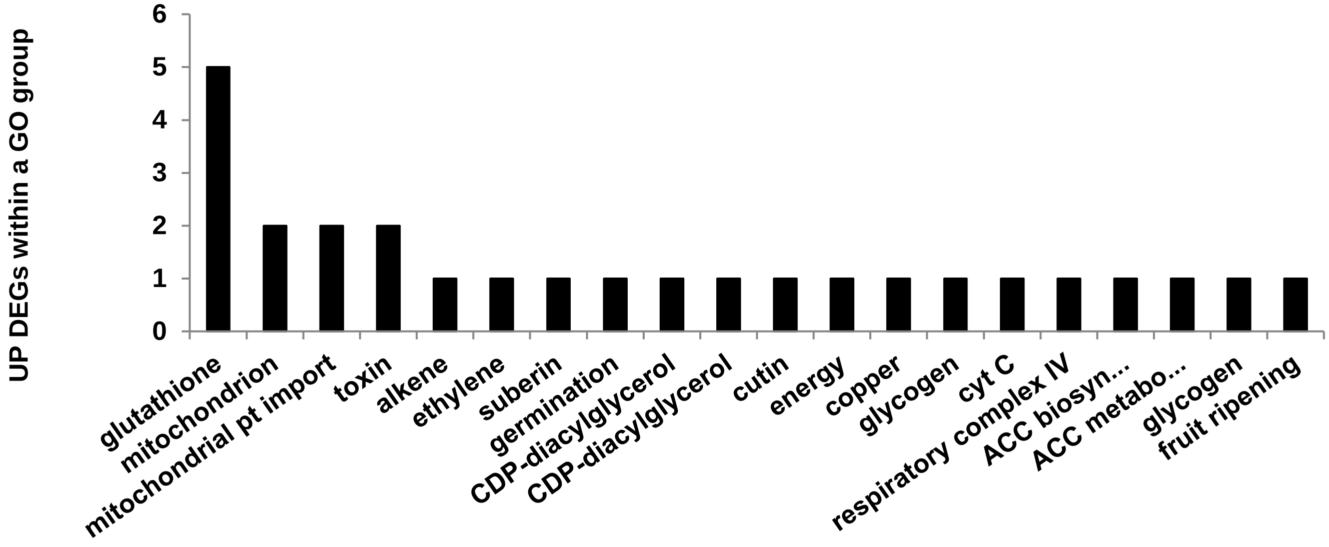


B

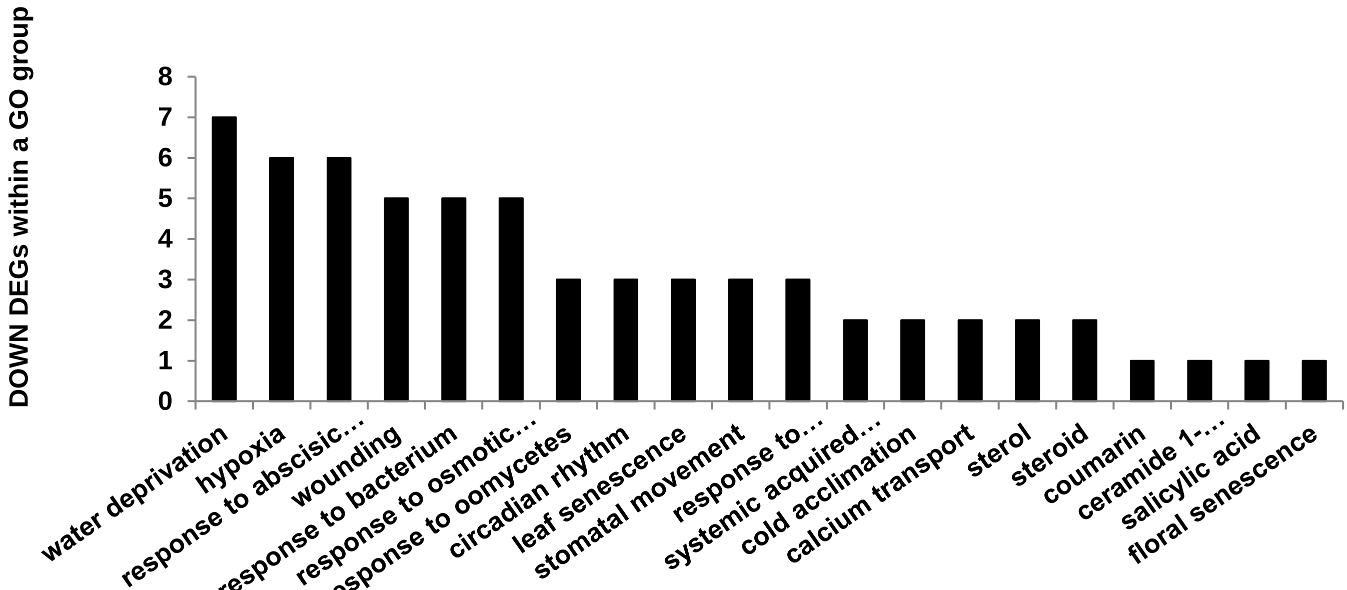


**Suppl Fig. 2**

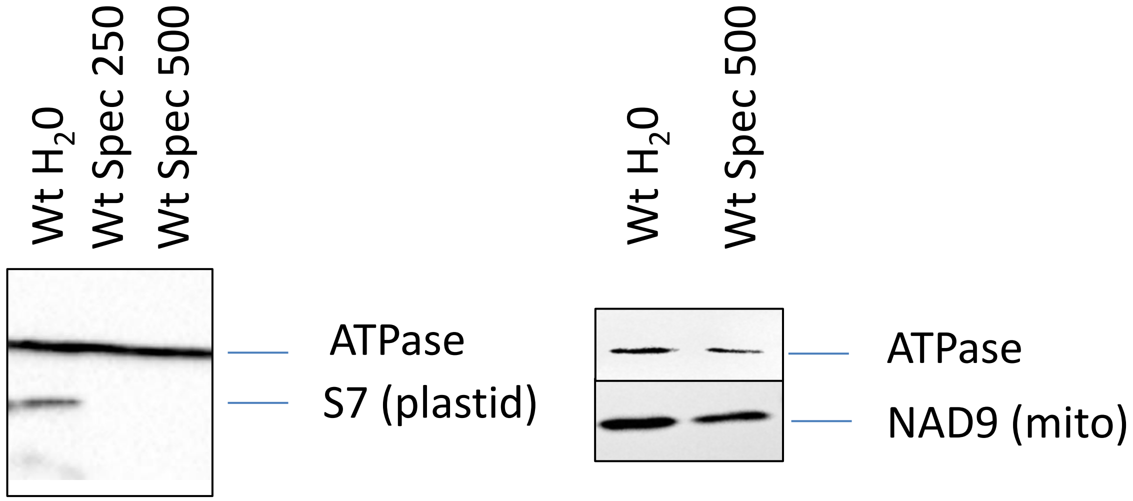

Wt $H_2O$
Wt Spec 250
Wt Spec 500
ATPase
S7 (plastid)
Wt $H_2O$
Wt Spec 500
ATPase
NAD9 (mito)


**Suppl Fig. 3**

A

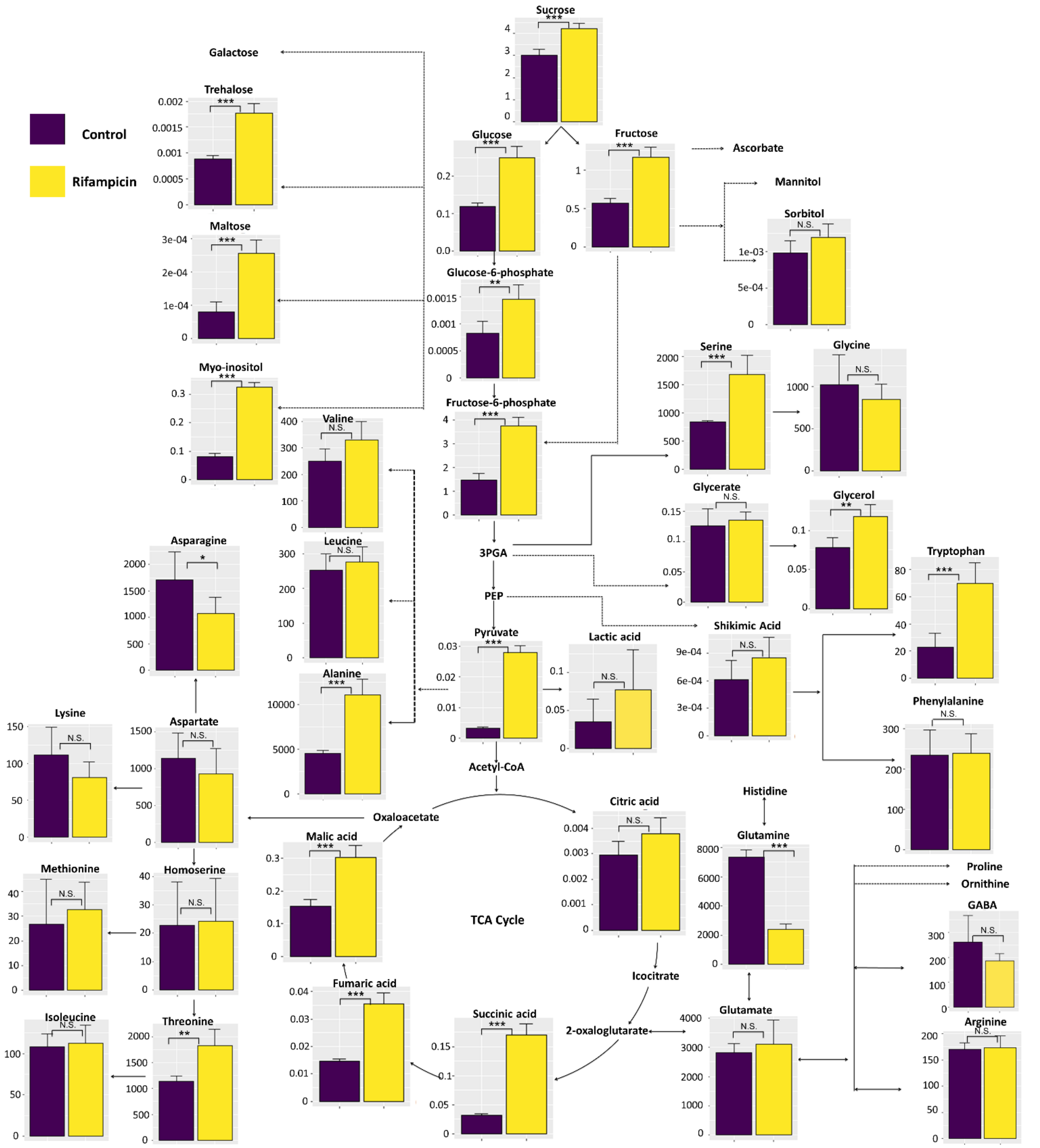


B

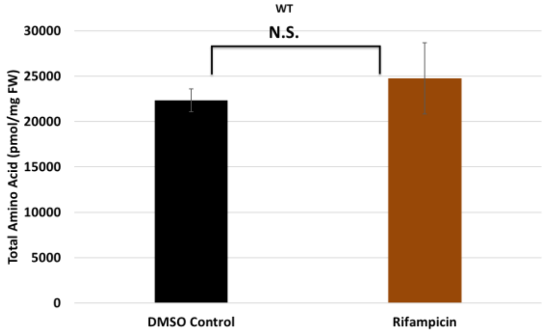


Suppl Fig. 4

A

| Gene | WT RIF/WT log2 FC | BH adj p-value | Protein | Signalling pathway |
|---|---|---|---|---|
| at2g20570 | 0,0 | 0,99 | GLK1 | PRG |
| at1g29910 | -0,4 | 0,76 | CAB3 | PRG |
| at2g05070 | -1,0 | 0,20 | LHCB2.2 | PRG |
| at5g38420 | -0,6 | 0,20 | RBCS-2B | PRG |
| at1g76100 | 0,0 | 0,99 | PETE1 | PRG |
| at3g01500 | 0,4 | 0,35 | CA1 | PRG |
| at2g34430 | 0,1 | 0,93 | LHB1B1 | PRG |
| at3g27690 | -0,3 | 0,63 | LHCB2.4 | PRG |
| at5g54190 | 0,0 | 0,97 | PORA | PRG |

B

| Gene | WT RIF/WT log2 FC | BH adj p-value | Protein | Synthesis/ signaling |
|---|---|---|---|---|
| at1g02500 | -0,02 | 0,96 | SAM1 | synthesis |
| at4g01850 | -0,12 | 0,65 | SAM2 | synthesis |
| at3g61510 | 0,25 | 0,59 | ACS1 | synthesis |
| at1g01480 | 0,18 | 0,76 | ACS2 | synthesis |
| at2g22810 | 1,99 | 0,003 | ACS4 | synthesis |
| at5g65800 | -0,78 | 0,12 | ACS5 | synthesis |
| at4g11280 | -0,96 | 0,01 | ACS6 | synthesis |
| at4g26200 | 0,03 | 0,97 | ACS7 | synthesis |
| at4g37770 | 0,59 | 0,34 | ACS8 | synthesis |
| at3g49700 | -0,04 | 0,97 | ACS9 | synthesis |
| at4g08040 | -0,13 | 0,85 | ACS11 | synthesis |
| at2g19590 | -0,43 | 0,48 | ACO1 | synthesis |
| at1g62380 | -0,07 | 0,88 | AC02 | synthesis |
| at1g05010 | -0,56 | 0,24 | ACO4 | synthesis |
| at2g25450 | -1,05 | 0,02 | 2-oxoglutarate-dependent dioxygenase | signaling |
| at3g12900 | -2,32 | 0,04 | Rolling-leaf1 4, 2OG-Fe (II) oxygenase | signaling |
| at3g24500 | -1,15 | 0,01 | ethylene responsive transcription factor | signaling |
| at5g25810 | -0,97 | 0,02 | TINY, ERF/AP2 transcription factor | signaling |
| at5g51190 | -1,04 | 0,12 | AP2 domain-containing transcription factor | signaling |

C

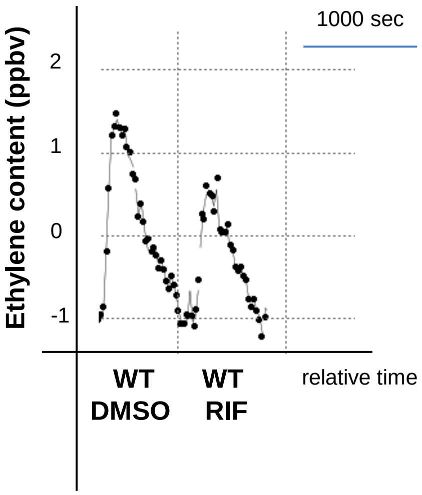


**Suppl Fig. 5**

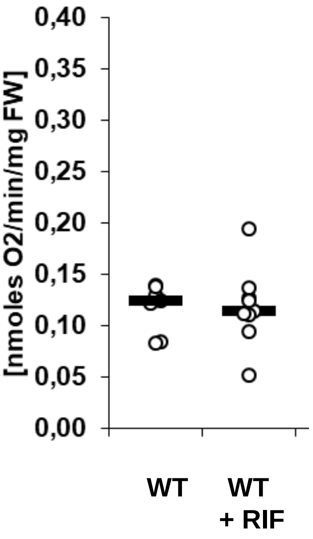


**Suppl Fig. 6**

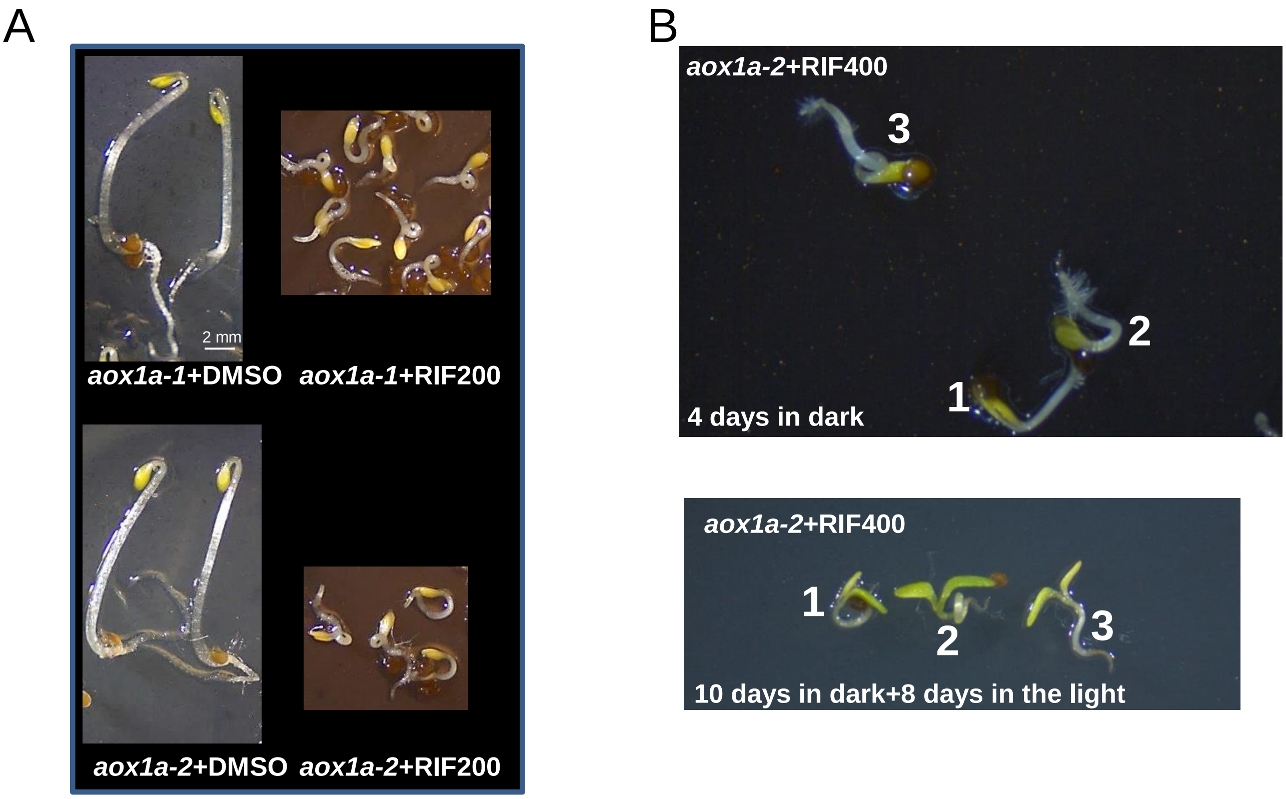
A
B
2 mm
aox1a-1+DMSO
aox1a-1+RIF200
aox1a-2+DMSO
aox1a-2+RIF200
aox1a-2+RIF400
3
2
1
4 days in dark
aox1a-2+RIF400
1
2
3
10 days in dark+8 days in the light

**Suppl Fig. 7**
